# Epidermal green autofluorescence intensity is significantly associated with the serum levels of multiple cytokines of LPS-exposed mice

**DOI:** 10.1101/823062

**Authors:** Yujia Li, Mingchao Zhang, Zhaoxia Yang, Weihai Ying

**Author notes:** Corresponding author Weihai Ying, Ph.D., Professor, School of Biomedical Engineering and Med-X Research Institute, Shanghai Jiao Tong University, 1954 Huashan Road, Shanghai, 200030, P.R. China. These two authors contributed equally to this work (Yujia Li, Mingchao Zhang).

## Abstract

Inflammation plays various crucial pathological and physiological roles, in which a number of pro-inflammatory and anti-inflammatory cytokines are important mediators. It is of great scientific and clinical significance to search for non-invasive approaches for evaluating cytokine levels in the blood. Our previous study reported that epidermal green autofluorescence (AF) intensity of LPS-exposed mice is highly correlated with LPS doses. In our current study, we determined if epidermal green AF intensity is associated with the serum levels of various cytokines in the LPS-exposed mice. We found that both epidermal green AF intensity and LPS doses are significantly associated with the serum levels of key cytokines including interleukin-1β (IL-1β), IL-6 and IL-10. Both epidermal green AF intensity and LPS doses are also significantly associated with the pro-inflammatory factors including IL-2, IL-12(p40), monocyte chemoattractant protein-1 (MCP-1), MIP-1α, MIP-1β, and regulated on activation, normal T cell expressed and secreted (RANTES/CCL5), as well as anti-inflammatory factors including IL-5 and granulocyte colony stimulating factor (G-CSF). Our findings have suggested that detection of epidermal green AF intensity may become first approach for non-invasive evaluation of certain cytokine levels in human body, which could profoundly enhance our capacity to evaluate inflammation levels for monitoring health state, disease state and therapeutic effects.

## Introduction

Inflammation is a critical pathological factor in multiple major diseases including cerebral ischemia (24), acute myocardial infarction (23), stable coronary artery disease (15), Parkinson’s disease (21) and lung cancer (19). A number of pro-inflammatory and anti-inflammatory cytokines are important mediators of inflammation (1-5,9,12,14,16,25,27). It is of great scientific and clinical significance to search for non-invasive approaches for evaluating cytokine levels in the blood.

Our previous study using mice exposed to lipopolysaccharide (LPS) has suggested that inflammation could be an important factor that can produce the increases in the epidermal green AF intensity, which may be used as the first biomarker for non-invasive and rapid evaluation of systemic inflammation (17, 29). However, it remains unclear if epidermal green AF intensity is also significantly associated with the levels of cytokines in the blood. If this association does exist, we may establish first non-invasive approach for evaluating the levels of cytokines in human body without the need of blood drawing, which could markedly enhance our capacity to evaluate the inflammation levels for monitoring health state, disease state and therapeutic effects.

In our current study, we determined if epidermal green AF intensity is associated with the serum levels of a number of cytokines in LPS-exposed mice. We found that both epidermal green AF intensity and LPS doses were significantly associated the serum levels of critical cytokines including IL-1β, IL-6 and IL-10. Both epidermal green AF intensity and LPS doses were also significantly associated with the pro-inflammatory factors including IL-2, IL-12(p40), MCP-1, MIP-1α, MIP-1β and RANTES, as well as anti-inflammatory factors including IL-5 and G-CSF.

## Materials and Methods

All chemicals were purchased from Sigma (St. Louis, MO, USA) except where noted.

### Animal Studies

Male C57BL/6Slac mice of SPF grade were purchased from SLRC Laboratory (Shanghai, China). All of the animal protocols were approved by the Animal Study Committee of the School of Biomedical Engineering, Shanghai Jiao Tong University.

### Administration of LPS

Male C57BL/6Slac mice with the weight of 20 - 25 g were administered with 0.5 or 1 mg/kg LPS with intraperitoneal (i.p.) injection. The stock solution of LPS with the final concentration of 0.2 mg/ml was made by dissolving LPS in PBS. The mice were administered with these doses of LPS every 24 hrs for three days.

### Imaging of epidermal AF

As described previously (13), the epidermal AF of the ears of the mice was imaged under a two-photon fluorescence microscope (A1 plus, Nikon Instech Co., Ltd., Tokyo, Japan), with the excitation wavelength of 488 nm and the emission wavelength of 500 - 530 nm. The AF was quantified as follows: Sixteen spots with the size of approximately 100 X 100 μm^2^ on the scanned images were selected randomly. After the average AF intensities of each layer were calculated, the sum of the average AF intensities of all layers of each spot was calculated, which is defined as the AF intensity of each spot.

### Determination of cytokine levels

Cytokine concentrations in the serum of mice were detected using a Bio-Plex Pro assay kit (Bio-Rad Laboratories). A total of 50 μl of antibody-conjugated beads was added to the assay plate. After the samples were diluted 1:4, 50 μl of diluted samples, standards, the blank, and the controls was added to the plate. The plate was incubated in the dark at room temperature (RT) with the shaking speed at 300 rpm for 30 min. After three washes with 100 μl wash buffer, a total of 25 μl biotinylated antibody was added to the plate. The plate was incubated in the dark at RT with shaking speed at 300 rpm for 30 min, which was washed 3 times with 100 μl washing buffer. After a total of 50 μl streptavidin-phycoerythrin (PE) was added to the plate, the plate was incubated in the dark at RT with shaking speed at 300 rpm for 10 min. After three washes, the plate was read using a Bio-Plex protein array reader. Bio-Plex Manager 6.0 software was used for the data acquisition and analysis of the assay.

### Statistical analyses

All data are presented as mean ± SEM. SPSS software was used to determine if distribution of the serum concentrations of each cytokine (in pg/ml) was in normal-distribution or non-normal distribution. An analysis showing *P* > 0.05 indicates the distribution was in normal-distribution, while an analysis showing *P* < 0.05 indicates the distribution was in non-normal distribution. For the cytokine concentrations that did not show normal-distribution, Log-transformation was conducted (10,22). Subsequently, SPSS software was used to conduct the linear regression between the cytokine concentrations or the Log-transformed cytokine concentrations and the LPS doses or the epidermal green AF intensity. The analyses generated *P* values and r values, respectively, which denote significance of linear trend and correlation coefficient, respectively. *P* values less than 0.05 were considered statistically significant.

## Results

### 1. Both epidermal green AF intensity and LPS doses were significantly associated the serum levels of IL-1β, IL-6 and IL-10 in LPS-exposed C57BL/6Slac mice

Since Interlukin-1β (IL-1β), IL-6 and IL-10 are key cytokines that regulate inflammatory processes (3,9,11,14,25,27), our study determined the relationship between LPS doses and the serum levels of these critical cytokines of the mice exposed to 0.5- or 1 mg / kg LPS three days after the LPS administration. Since none of the serum level of IL-1β, IL-6 or IL-10 was in normal distribution, Log transformation of the cytokine levels was conducted for the data association analyses. We found that LPS doses were significantly associated with Log-transformed serum level of IL-1β - a pro-inflammatory factor that plays a key a role in host responses to endogenous and exogenous noxious stimuli (9) (Fig. 1A). Epidermal green AF intensity was also significantly associated with Log-transformed serum level of IL-1β (Fig. 1B). We further found that both LPS doses and the epidermal green AF intensity were significantly associated with Log-transformed serum level of IL-6 - a cytokine that plays significant roles in host defense through stimulation of acute phase responses, immune reactions and hematopoiesis (27) (Fig. 2A and Fig. 2B). Both LPS doses and the AF intensity were also significantly associated with the Log-transformed serum level of IL-10 - a pleiotropic cytokine that is involved in the pathogenesis and development of autoimmune diseases and cancer (14) (Fig. 3A and Fig. 3B).

**Fig. 1.**
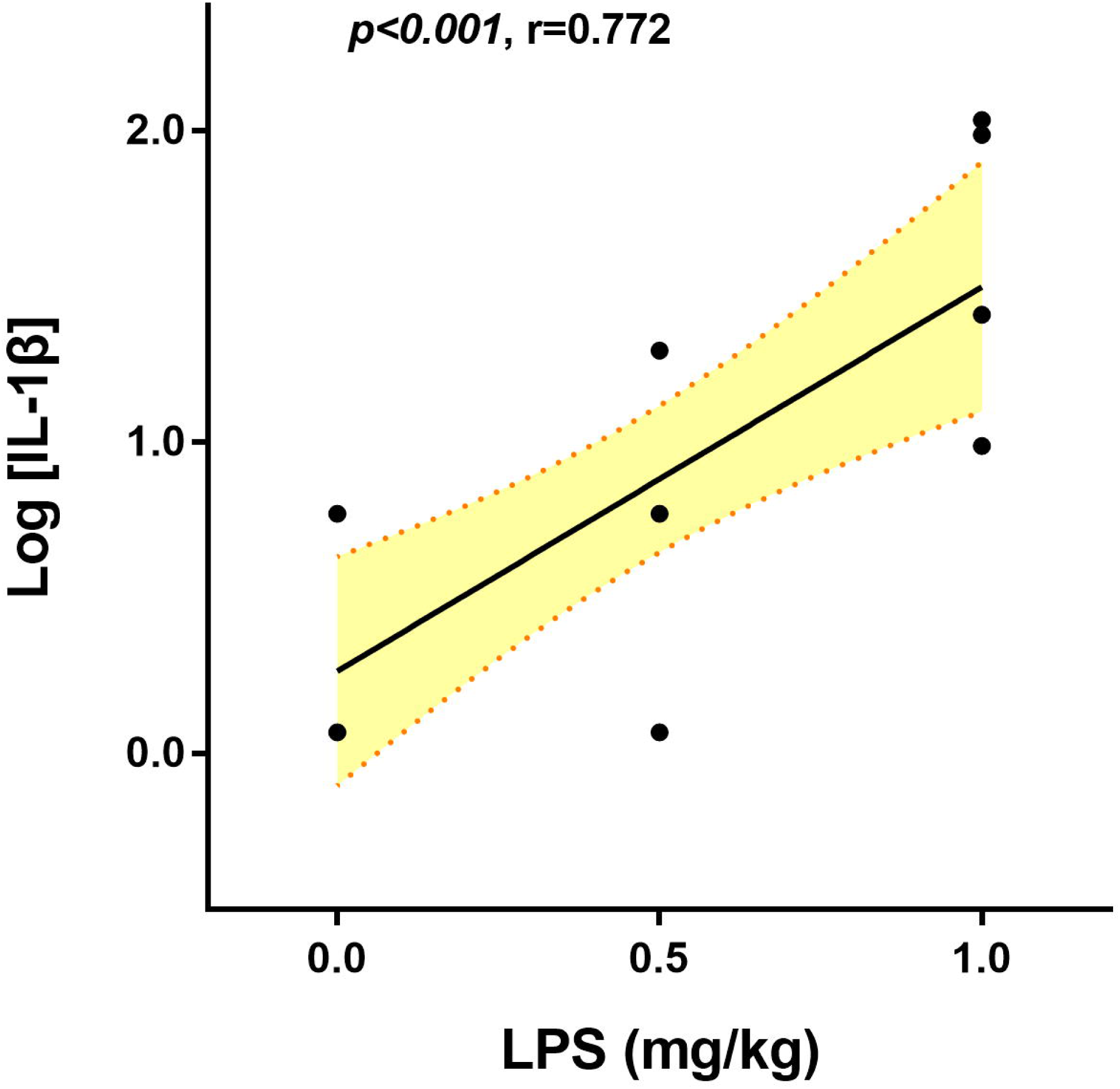
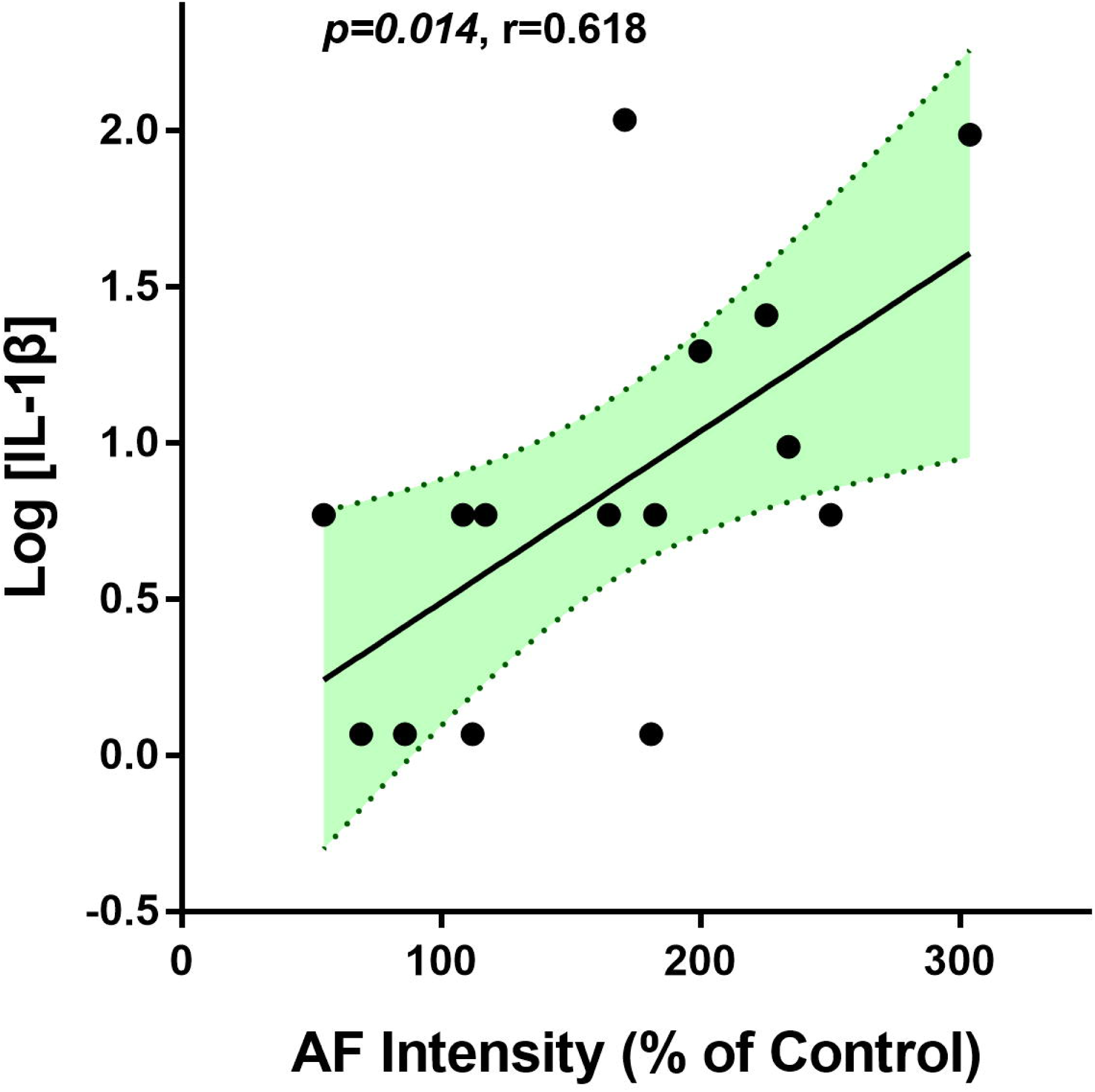
Both LPS doses and the epidermal green AF intensity were significantly associated the Log-transformed serum level of IL-1β in LPS-exposed C57BL/6Slac mice. Both LPS doses (A) and the epidermal green AF intensity (B) were significantly associated the serum level of IL-1β in LPS-exposed C57BL/6Slac mice. Each day the mice were i.p. administered with 0.5 or 1 mg / kg LPS. Three days after the first LPS administration, the serum level of IL-1β was determined. The serum IL-1β level (in pg/ml) was log-transformed. There were 4 - 6 mice in each group. The area in green shadow represents 95% confidence interval around the fitted line at the population level.

**Fig. 2.**
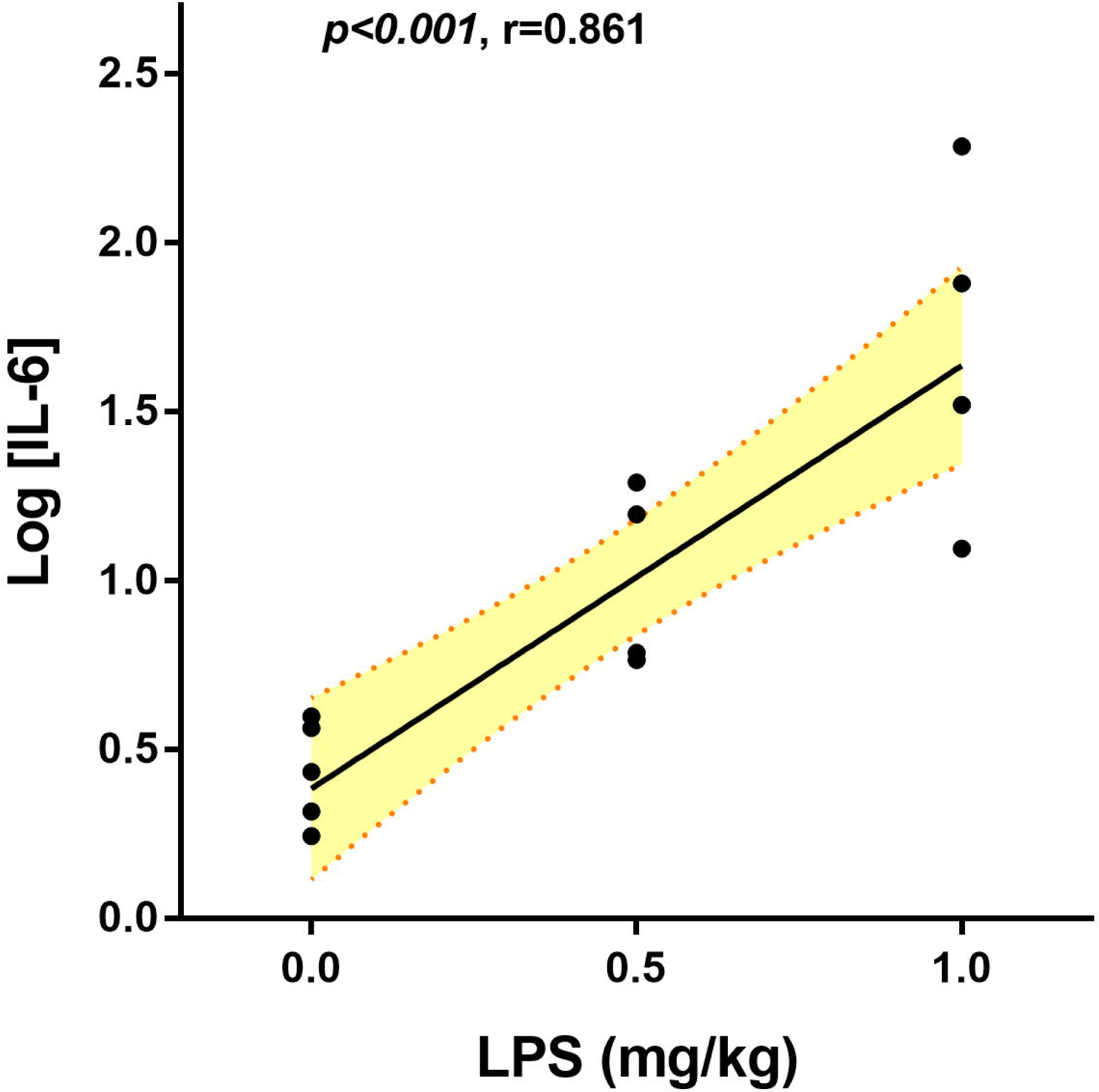
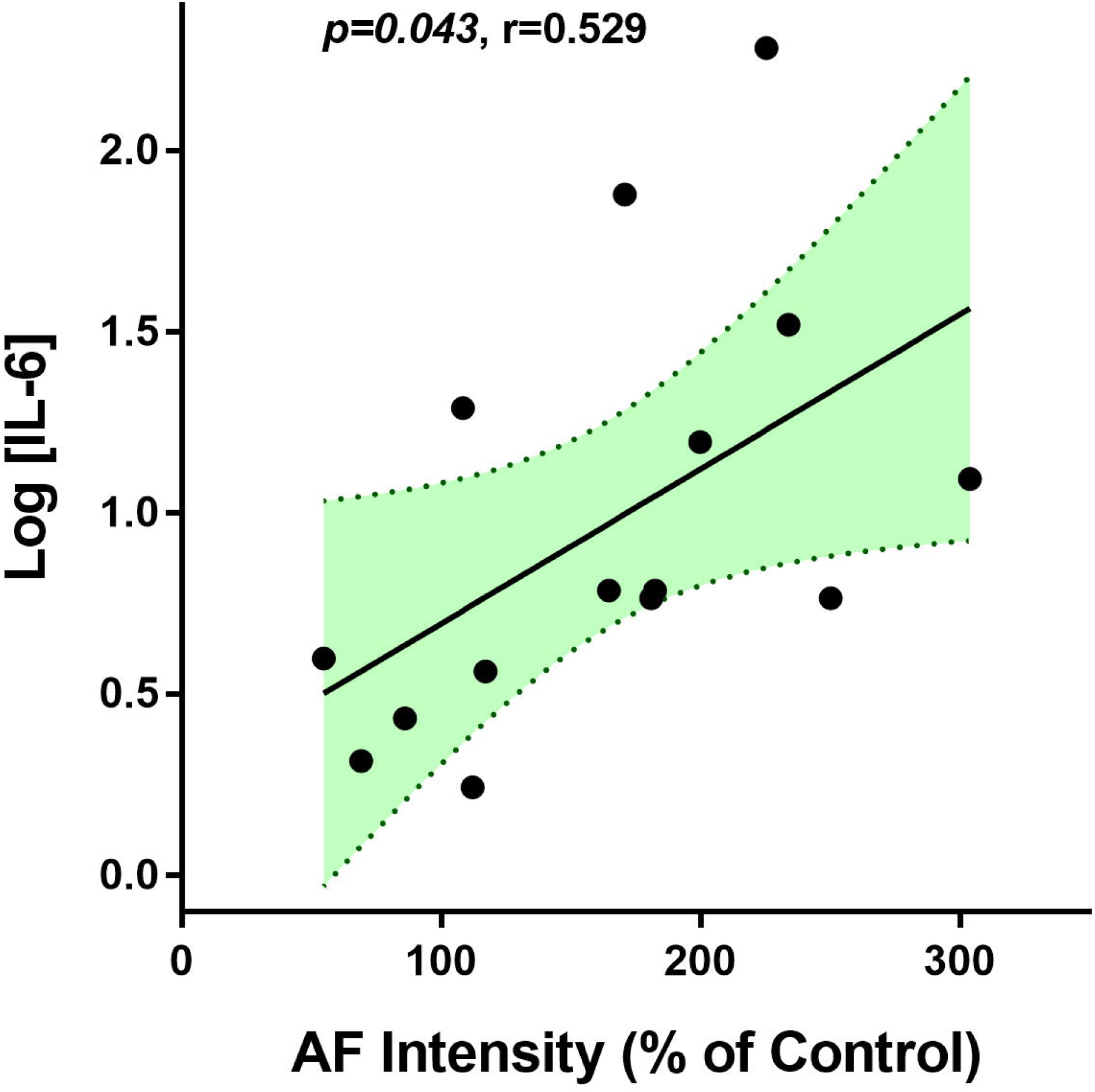
Both LPS doses and the epidermal green AF intensity were significantly associated the Log-transformed serum level of IL-6 in LPS-exposed C57BL/6Slac mice. Both LPS doses (A) and the epidermal green AF intensity (B) were significantly associated the serum level of IL-1β in LPS-exposed C57BL/6Slac mice. Each day the mice were i.p. administered with 0.5 or 1 mg / kg LPS. Three days after the first LPS administration, the serum level of IL-6 was determined. The serum IL-6 level (in pg/ml) was log-transformed. There were 4 - 6 mice in each group. The area in green shadow represents 95% confidence interval around the fitted line at the population level.

**Fig. 3.**
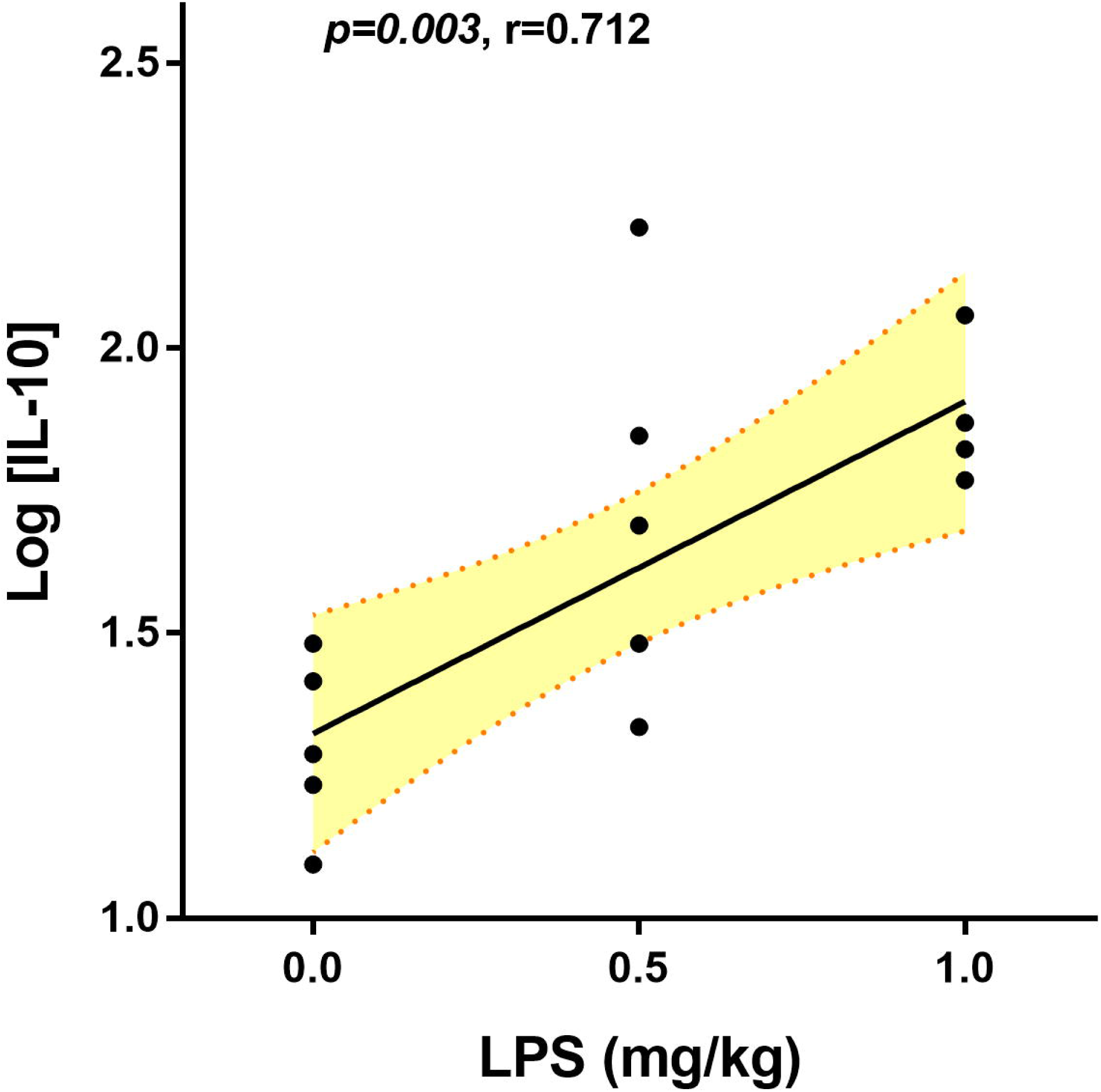
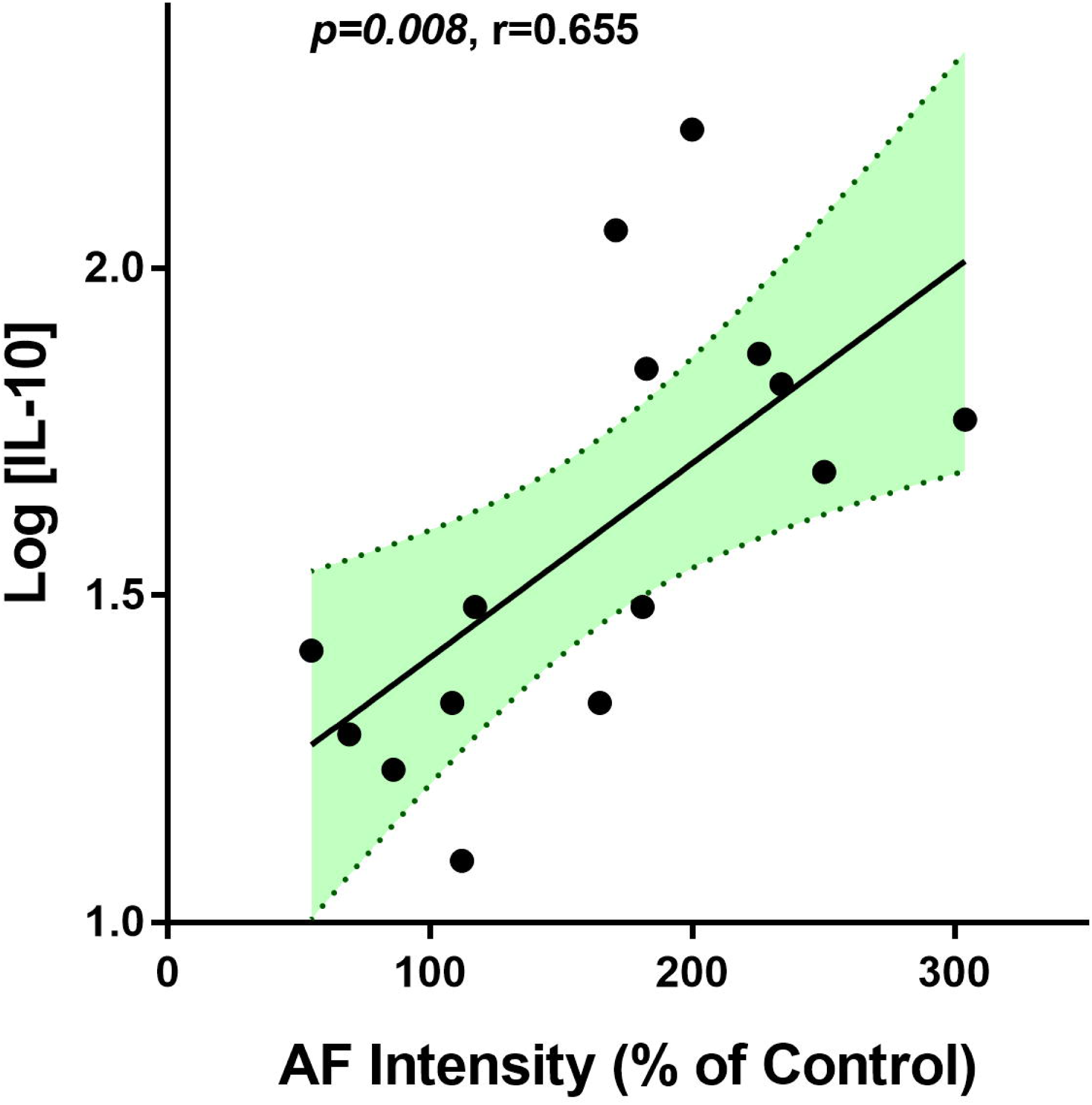
Both LPS doses and the epidermal green AF intensity were significantly associated the Log-transformed serum level of IL-10 in LPS-exposed C57BL/6Slac mice. Both LPS doses (A) and the epidermal green AF intensity (B) were significantly associated the serum level of IL-10 in LPS-exposed C57BL/6Slac mice. Each day the mice were i.p. administered with 0.5 or 1 mg / kg LPS. Three days after the first LPS administration, the serum level of IL-6 was determined. The serum IL-10 level (in pg/ml) was log-transformed. There were 4 - 6 mice in each group. The area in green shadow represents 95% confidence interval around the fitted line at the population level.

### 2. Both epidermal green AF intensity and LPS doses were significantly associated the serum levels of the pro-inflammatory factors including IL-2, IL-12(p40), MCP, MIP-1α, MIP-1β and RANTES/CCL5 in LPS-exposed C57BL/6Slac mice

IL-2 is a key cytokine in regulation of immune activation and tolerance by its effects on cytotoxic effector lymphocytes and CD4^+^ T regulatory (Treg) cells, respectively (2). IL-12(p40) is a common subunit shared by IL-12 and IL-23, which plays critical pathological roles in tumor growth, inflammatory bowel disease and psoriasis (5). MCP-1 is a chemokine that regulates monocyte chemotaxis and T-lymphocyte differentiation by binding to the CC chemokine receptor 2 (CCR2), which is a critical pathological factor in atherosclerosis, cancer and inflammatory diseases (1). Since none of the serum level of IL-2, IL-12(p40) or MCP-1 was in normal distribution, Log transformation was conducted for the data association analyses. We found that both LPS doses and the epidermal green AF intensity were significantly associated with Log-transformed serum level of IL-2 (Fig. 4A and Fig. 4B), IL-12(p40) (Fig. 5A and Fig. 5B) and MCP-1 (Fig. 6A and Fig. 6B).

**Fig. 4.**
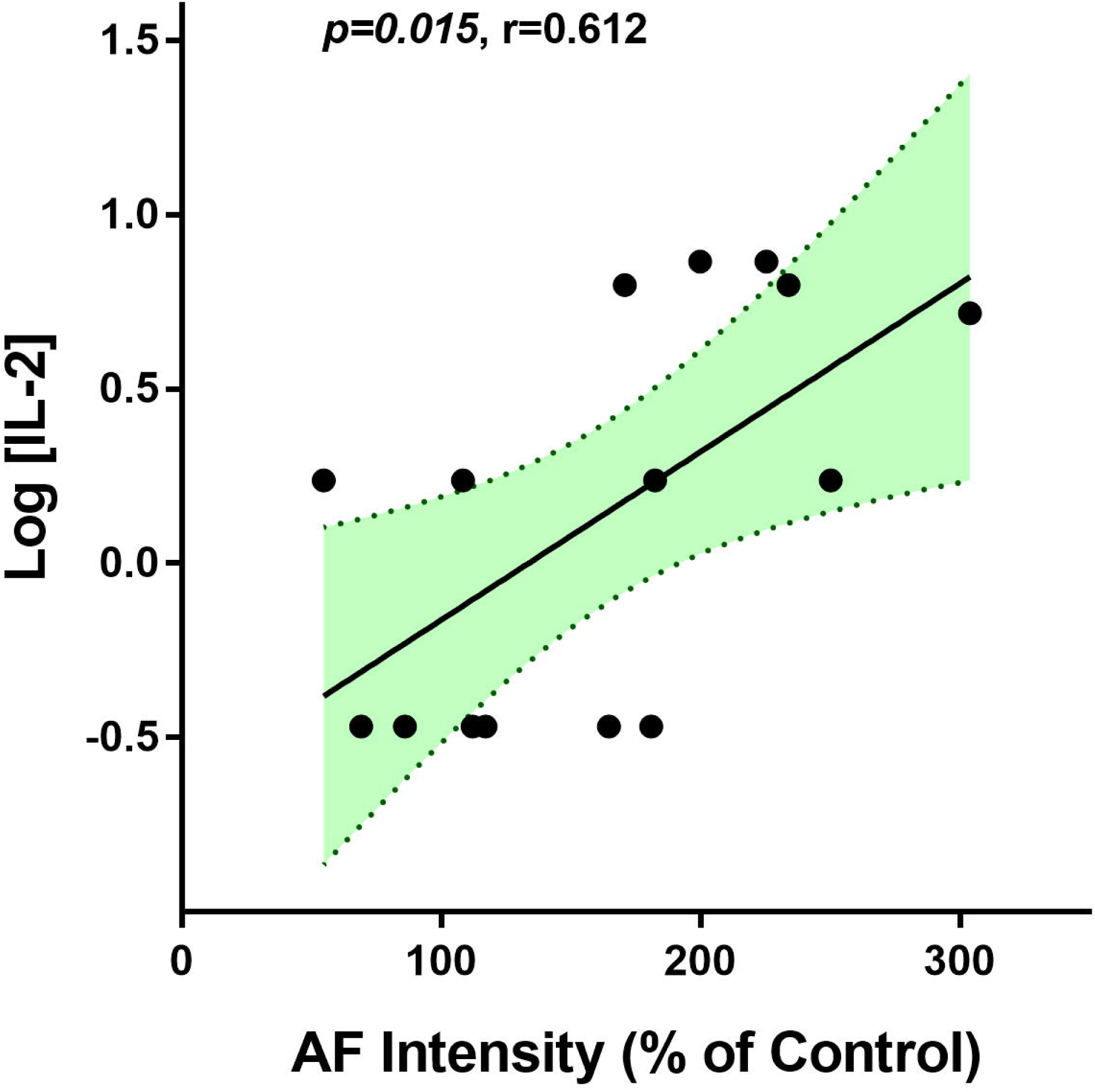
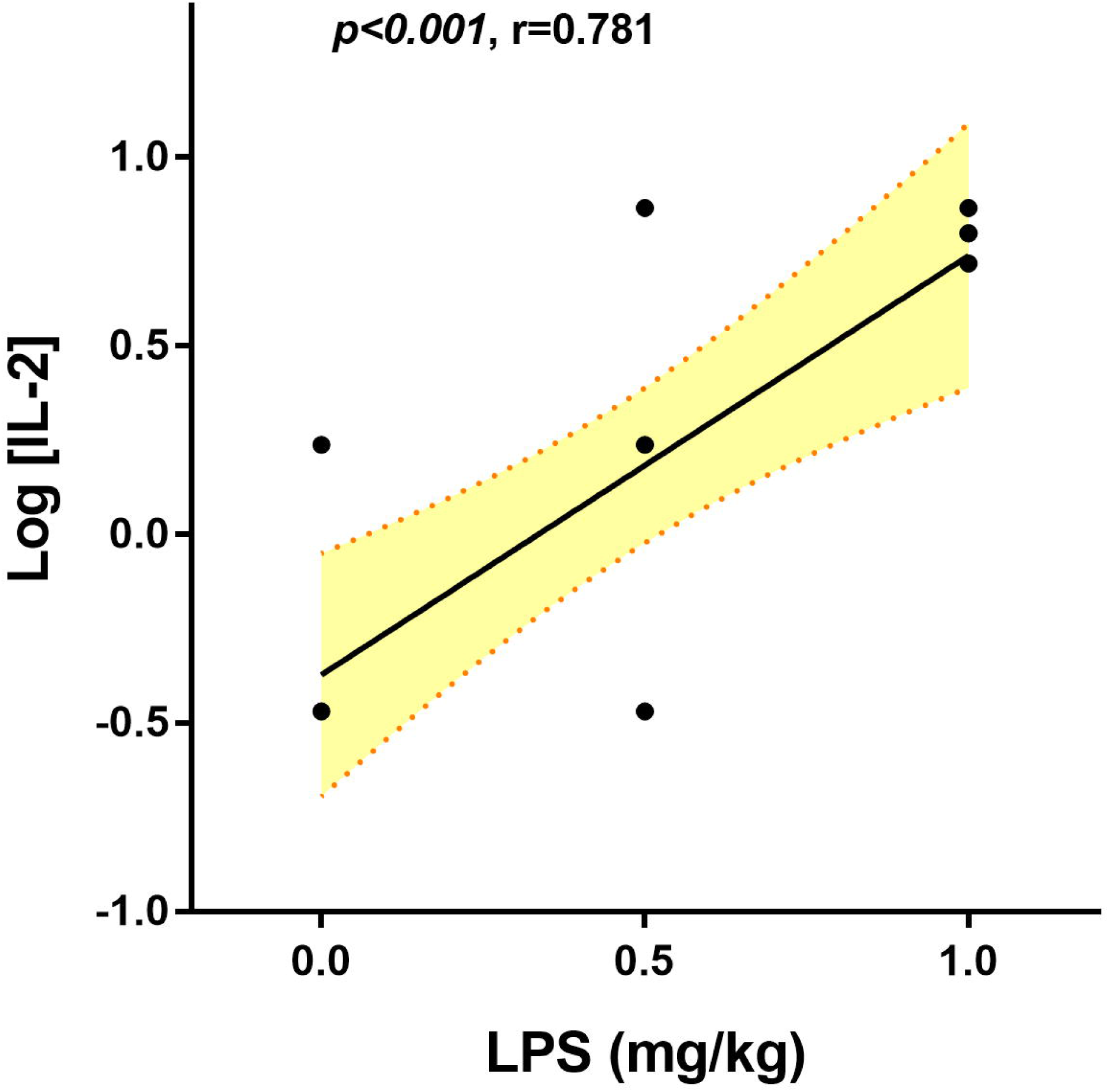
Both LPS doses and the epidermal green AF intensity were significantly associated the Log-transformed serum level of IL-2 in LPS-exposed C57BL/6Slac mice. Both LPS doses (A) and the epidermal green AF intensity (B) were significantly associated the serum level of IL-2 in LPS-exposed C57BL/6Slac mice. Each day the mice were i.p. administered with 0.5 or 1 mg / kg LPS. Three days after the first LPS administration, the serum level of IL-6 was determined. The serum IL-2 level (in pg/ml) was log-transformed. There were 4 - 6 mice in each group. The area in green shadow represents 95% confidence interval around the fitted line at the population level.

**Fig. 5.**
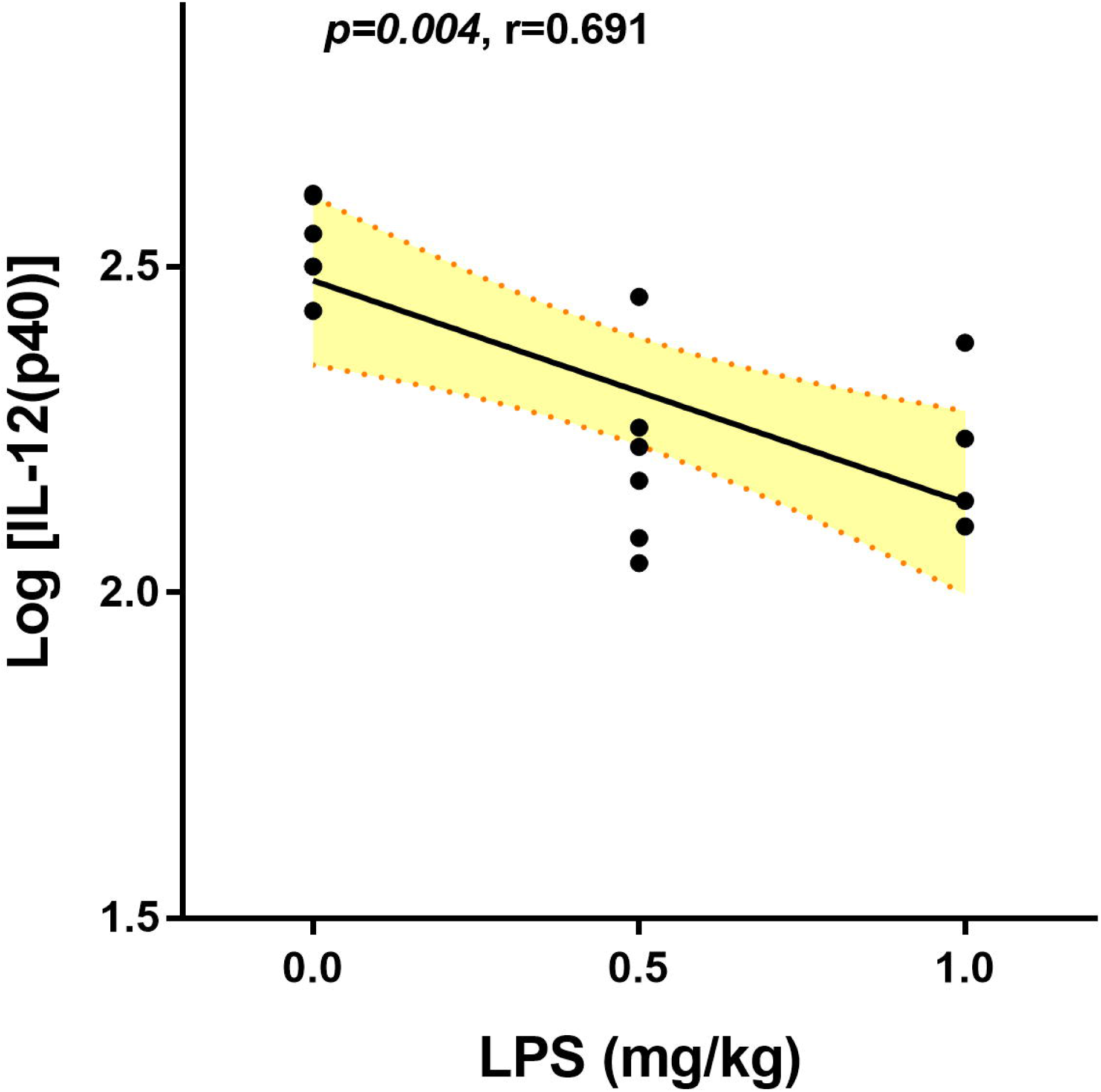
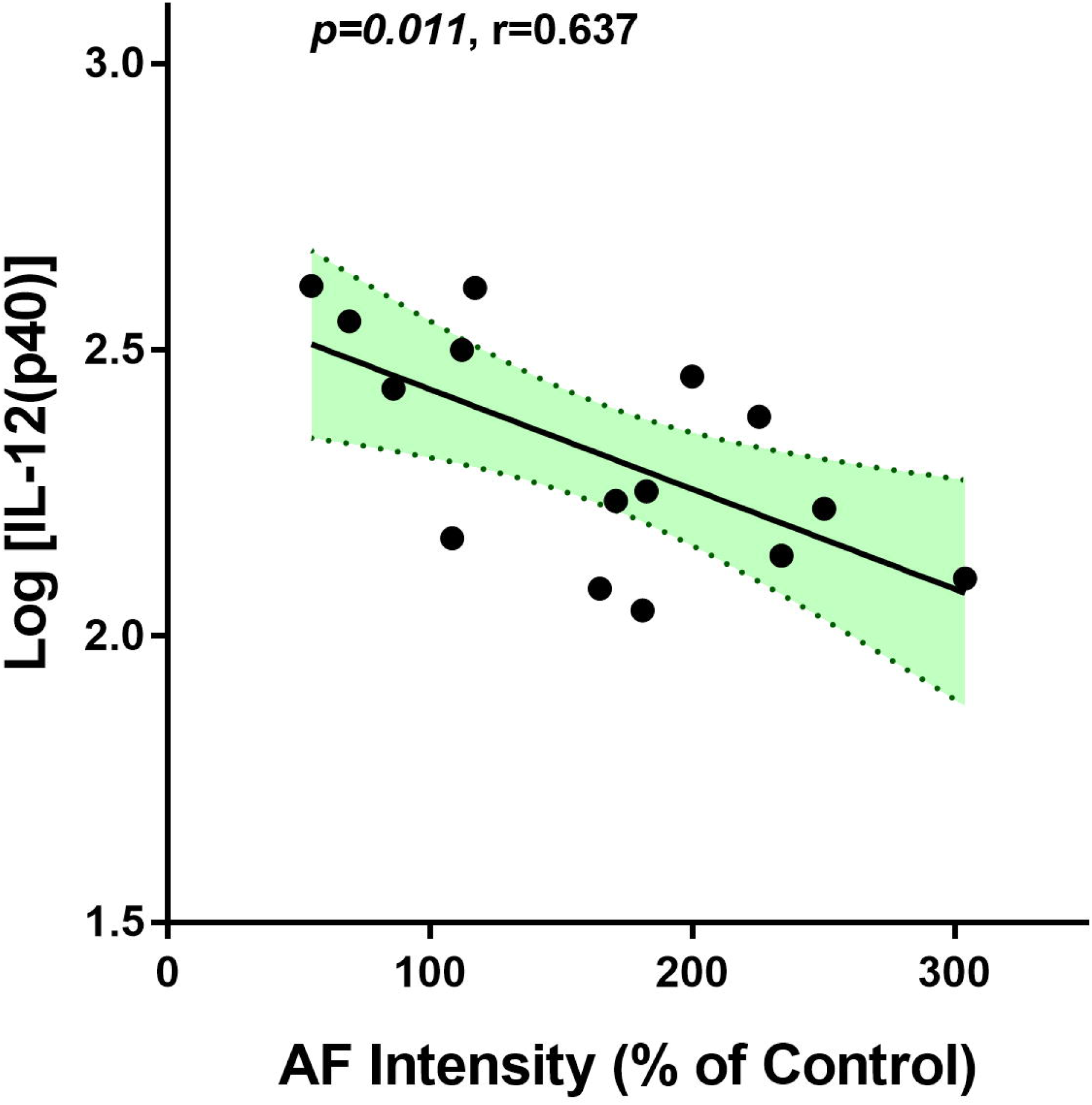
Both LPS doses and the epidermal green AF intensity were significantly associated the Log-transformed serum level of IL-12(p40) in LPS-exposed C57BL/6Slac mice. Both LPS doses (A) and the epidermal green AF intensity (B) were significantly associated the serum level of IL-12(p40) in LPS-exposed C57BL/6Slac mice. Each day the mice were i.p. administered with 0.5 or 1 mg / kg LPS. Three days after the first LPS administration, the serum level of IL-12(p40) was determined. The serum IL-12(p40) level (in pg/ml) was log-transformed. There were 4 - 6 mice in each group. The area in green shadow represents 95% confidence interval around the fitted line at the population level.

**Fig. 6.**
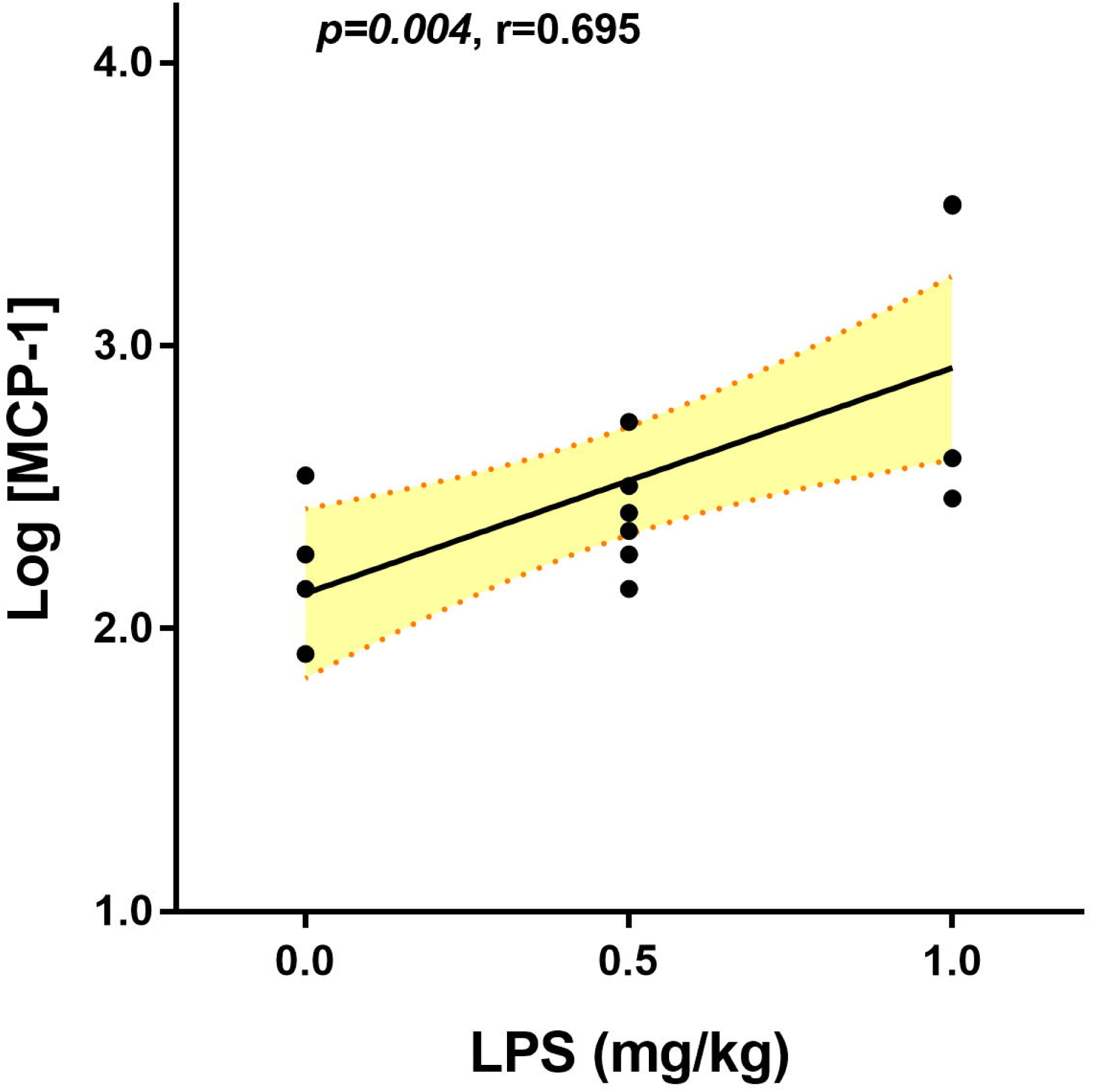
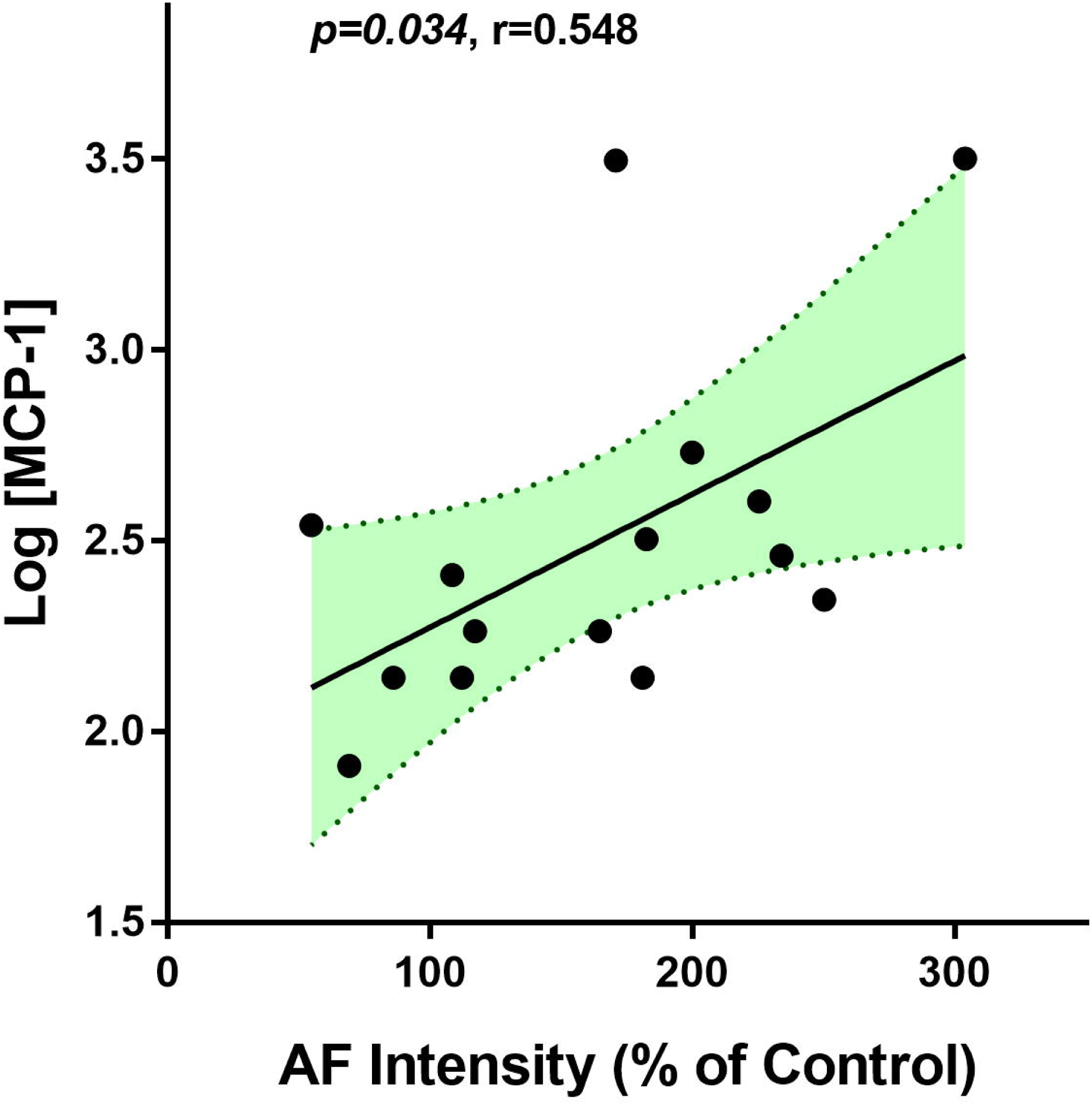
Both LPS doses and the epidermal green AF intensity were significantly associated the Log-transformed serum level of MCP-1 in LPS-exposed C57BL/6Slac mice. Both LPS doses (A) and the epidermal green AF intensity (B) were significantly associated the serum level of MCP-1 in LPS-exposed C57BL/6Slac mice. Each day the mice were i.p. administered with 0.5 or 1 mg / kg LPS. Three days after the first LPS administration, the serum level of MCP-1 was determined. The serum MCP-1 level (in pg/ml) was log-transformed. There were 4 - 6 mice in each group. The area in green shadow represents 95% confidence interval around the fitted line at the population level.

Regulated on activation, normal T cell expressed and secreted (RANTES) is an important chemokine for migration and homing of effector and memory T cells during acute infections (4). Since the serum levels of MIP-1α, MIP-1β and RANTES were in normal distribution, no Log transformation was needed for the data association analyses. We found that both LPS doses and the epidermal green AF intensity were significantly associated with the serum level of MIP-1α (Fig. 7A and Fig. 7B), MIP-1β (Fig. 8A and Fig. 8B) or RANTES (Fig. 9A and Fig. 9B) in the LPS-exposed mice. The log-transformed serum level of IL-1α was significantly associated with LPS doses (*p* < 0.001) (Fig. 10A), while it was only marginally associated with the AF intensity (*p* = 0.050) (Fig. 10B).

**Fig. 7.**
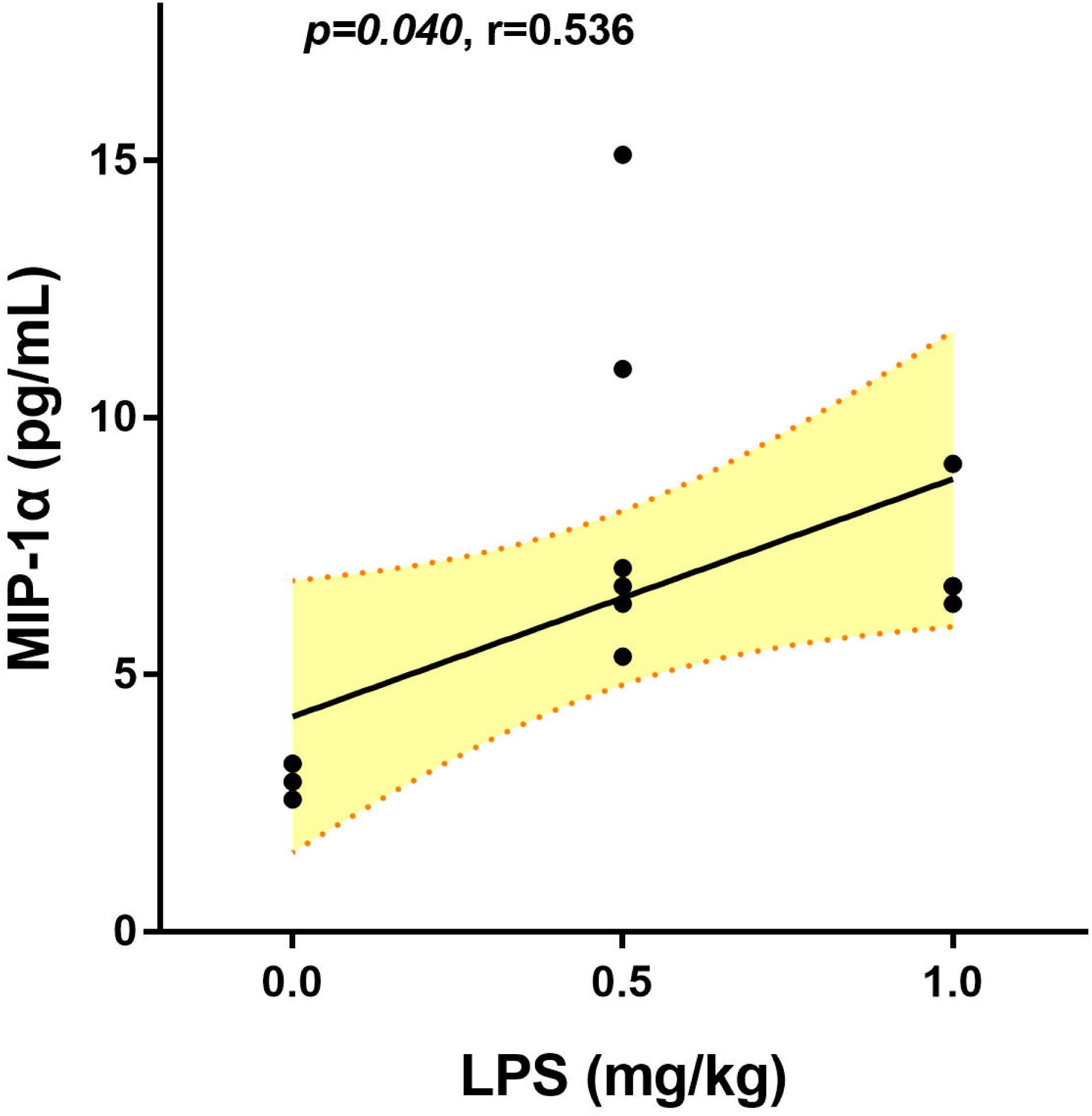
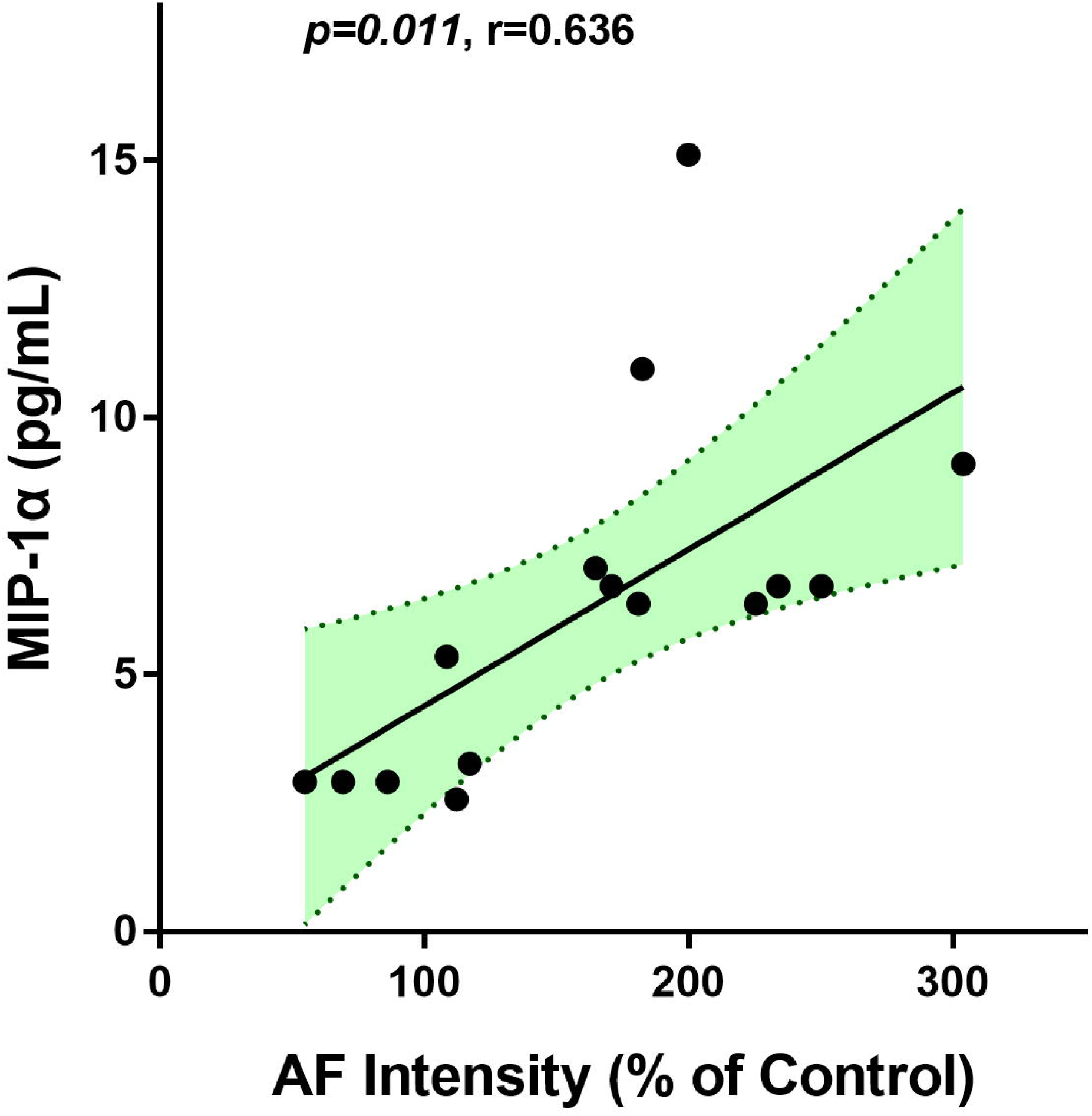
Both LPS doses and the epidermal green AF intensity were significantly associated the serum level of MIP-1α in LPS-exposed C57BL/6Slac mice. Both LPS doses (A) and the epidermal green AF intensity (B) were significantly associated the serum level of MIP-1α in LPS-exposed C57BL/6Slac mice. Each day the mice were i.p. administered with 0.5 or 1 mg / kg LPS. Three days after the first LPS administration, the serum level of the cytokines MIP-1α was determined. There were 4 - 6 mice in each group. The area in green shadow represents 95% confidence interval around the fitted line at the population level.

**Fig. 8.**
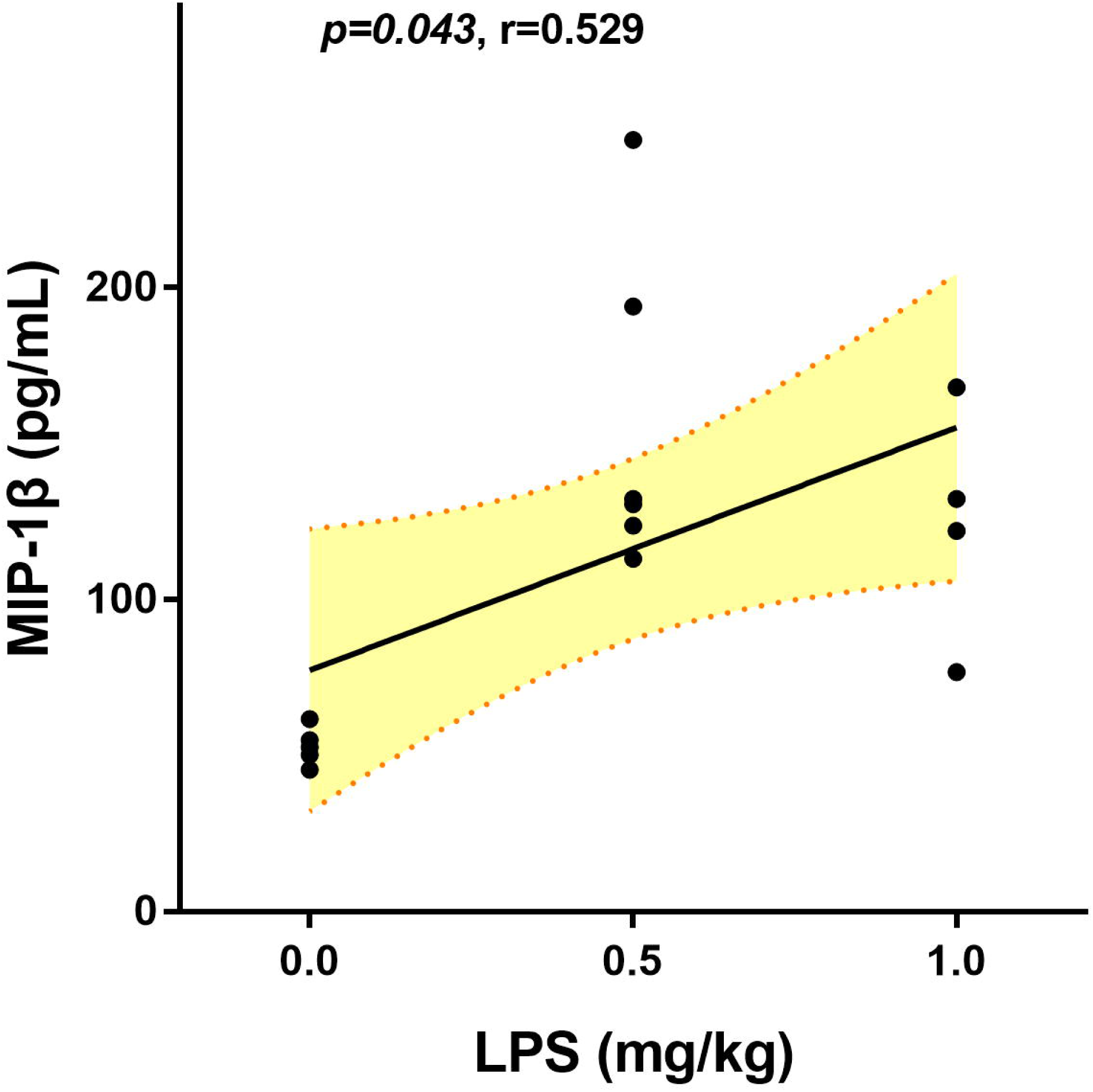
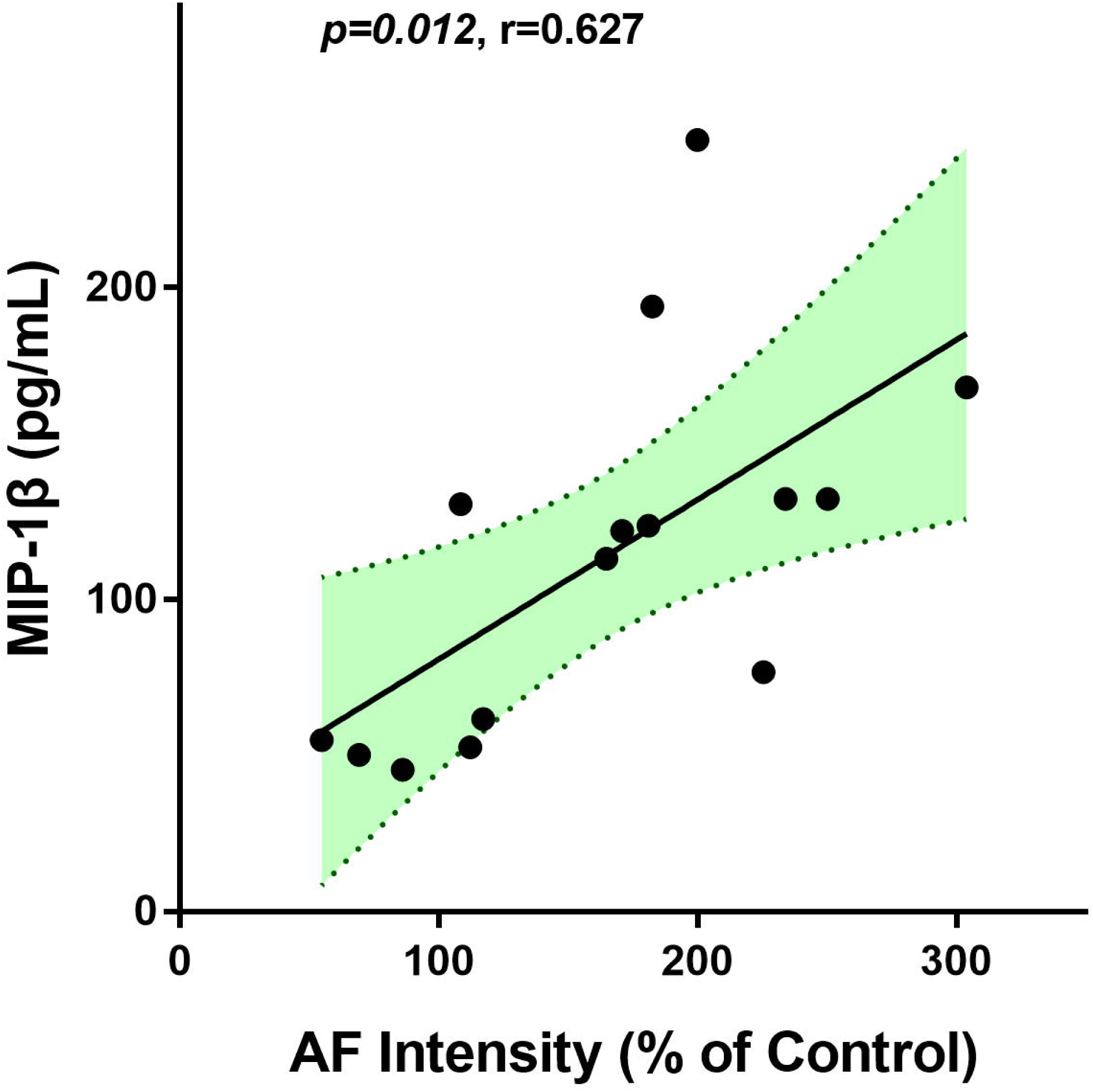
Both LPS doses and the epidermal green AF intensity were significantly associated the serum level of MIP-1β in LPS-exposed C57BL/6Slac mice. Both LPS doses (A) and the epidermal green AF intensity (B) were significantly associated the serum level of MIP-1β in LPS-exposed C57BL/6Slac mice. Each day the mice were i.p. administered with 0.5 or 1 mg / kg LPS. Three days after the first LPS administration, the serum level of the cytokines MIP-1β was determined. There were 4 - 6 mice in each group. The area in green shadow represents 95% confidence interval around the fitted line at the population level.

**Fig. 9.**
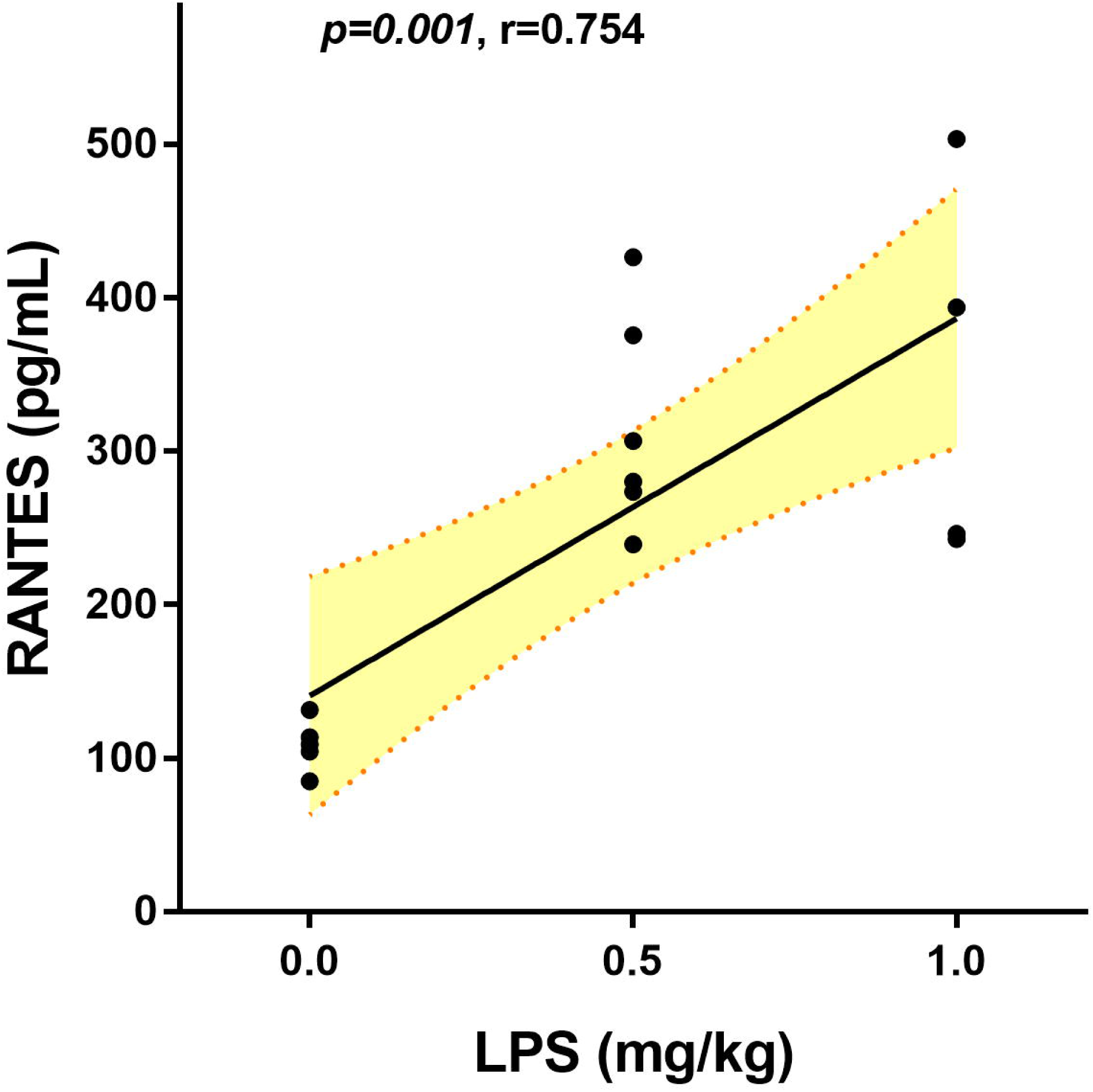
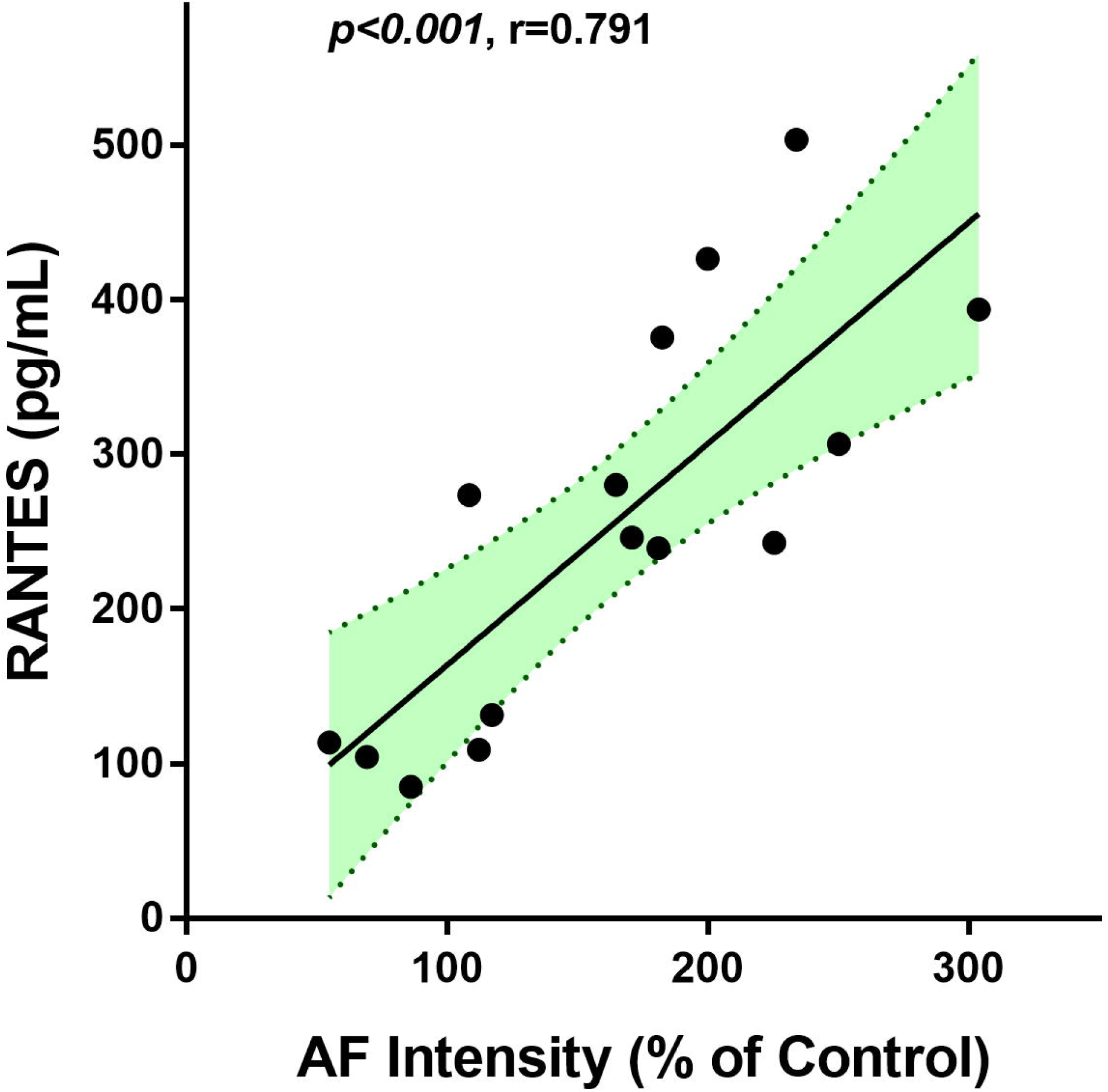
Both LPS doses and the epidermal green AF intensity were significantly associated the serum level of RANTES in LPS-exposed C57BL/6Slac mice. Both LPS doses (A) and the epidermal green AF intensity (B) were significantly associated the serum level of RANTES in LPS-exposed C57BL/6Slac mice. Each day the mice were i.p. administered with 0.5 or 1 mg / kg LPS. Three days after the first LPS administration, the serum level of RANTES was determined. There were 4 – 6 mice in each group. The area in green shadow represents 95% confidence interval around the fitted line at the population level.

**Fig. 10.**
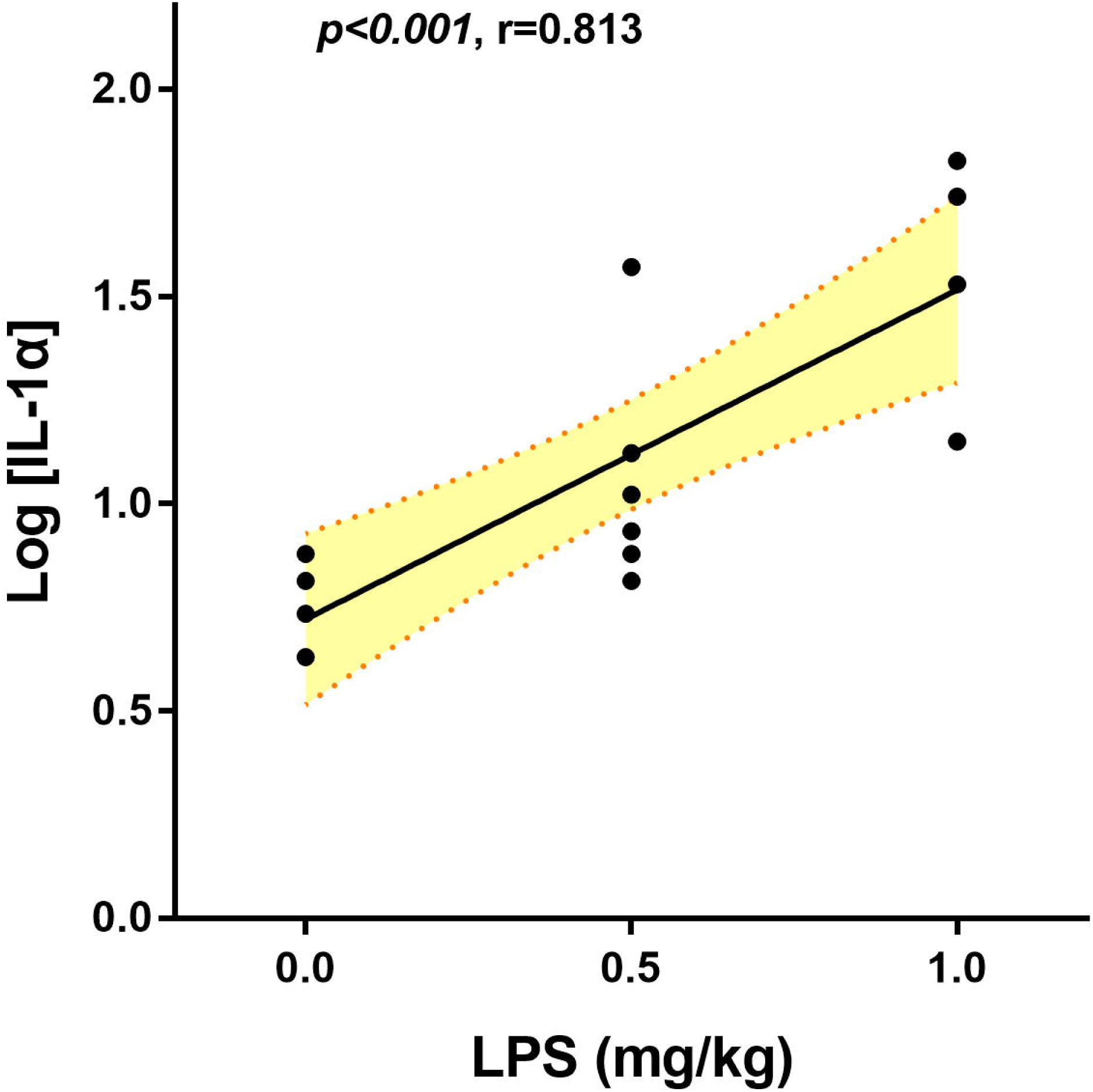
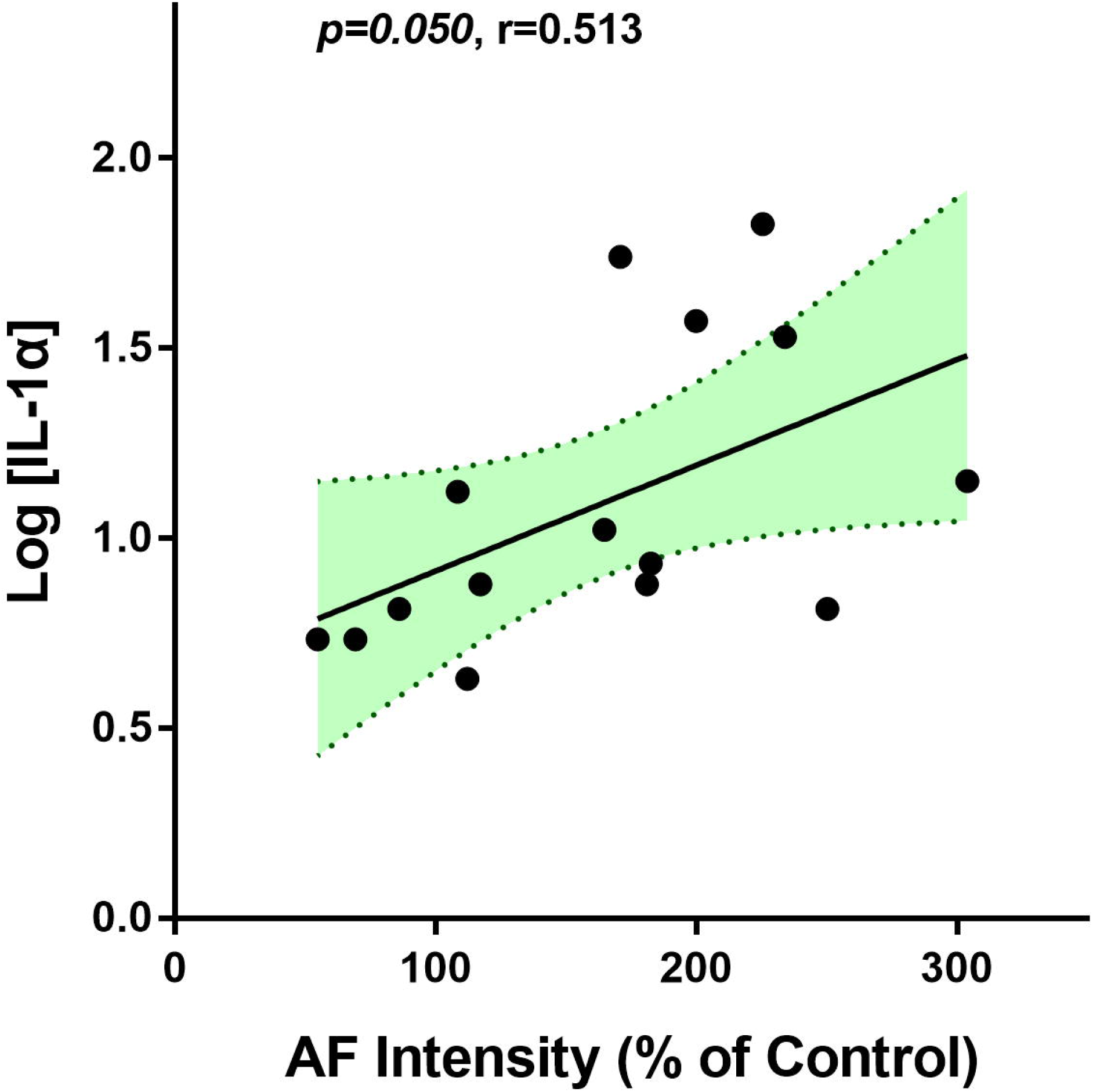
LPS doses was significantly associated the Log-transformed serum level of IL-1α in LPS-exposed C57BL/6Slac mice. (A) LPS doses was significantly associated the Log-transformed serum level of IL-1α in LPS-exposed C57BL/6Slac mice. (B) The epidermal green AF intensity was only marginally associated the Log-transformed serum level of IL-1α in LPS-exposed C57BL/6Slac mice. Each day the mice were i.p. administered with 0.5 or 1 mg / kg LPS. Three days after the first LPS administration, the serum level of IL-1α was determined. The serum IL-1α level (in pg/ml) was log-transformed. There were 4 - 6 mice in each group. The area in green shadow represents 95% confidence interval around the fitted line at the population level.

### 3. Both epidermal green AF intensity and LPS doses were significantly associated the serum levels of the anti-inflammatory factors including IL-5 and granulocyte colony stimulating factor (G-CSF) in LPS-exposed C57BL/6Slac mice

IL-5 plays a key role in the proliferation, maturation, activation, recruitment and survival of eosinophils (12). G-CSF is a hematopoietic growth factor that plays significant roles in the development of committed progenitor cells to neutrophils as well as the production, differentiation and function of granulocytes (16). Since none of the serum level of IL-5 or G-CSF was in normal distribution, Log transformation was conducted for the data association analyses. We found that both LPS doses and the epidermal green AF intensity were significantly associated with Log-transformed serum level of IL-5 (Fig. 11A and Fig. 11B) and G-CSF (Fig. 12A and Fig. 12B).

**Fig. 11.**
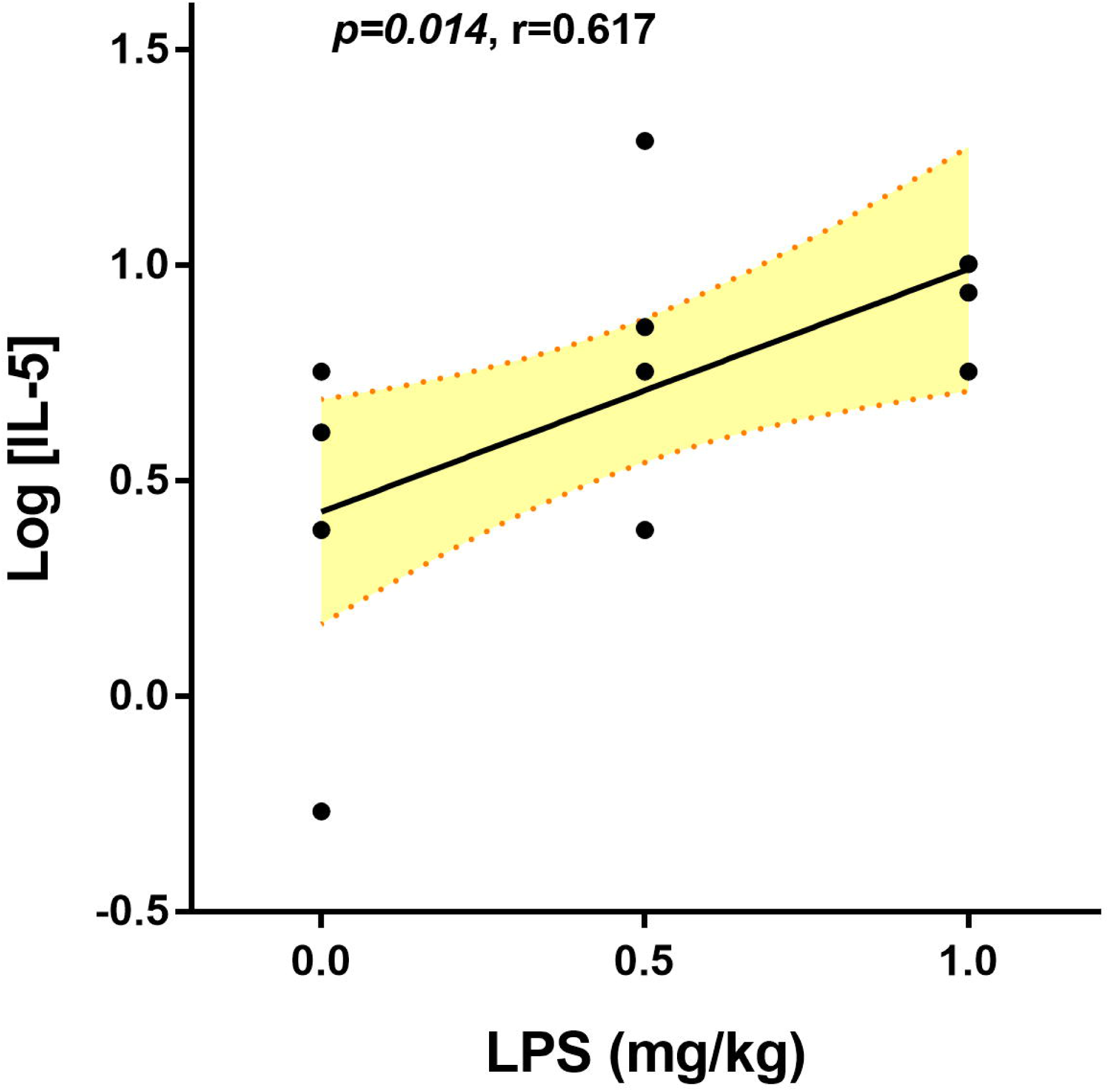
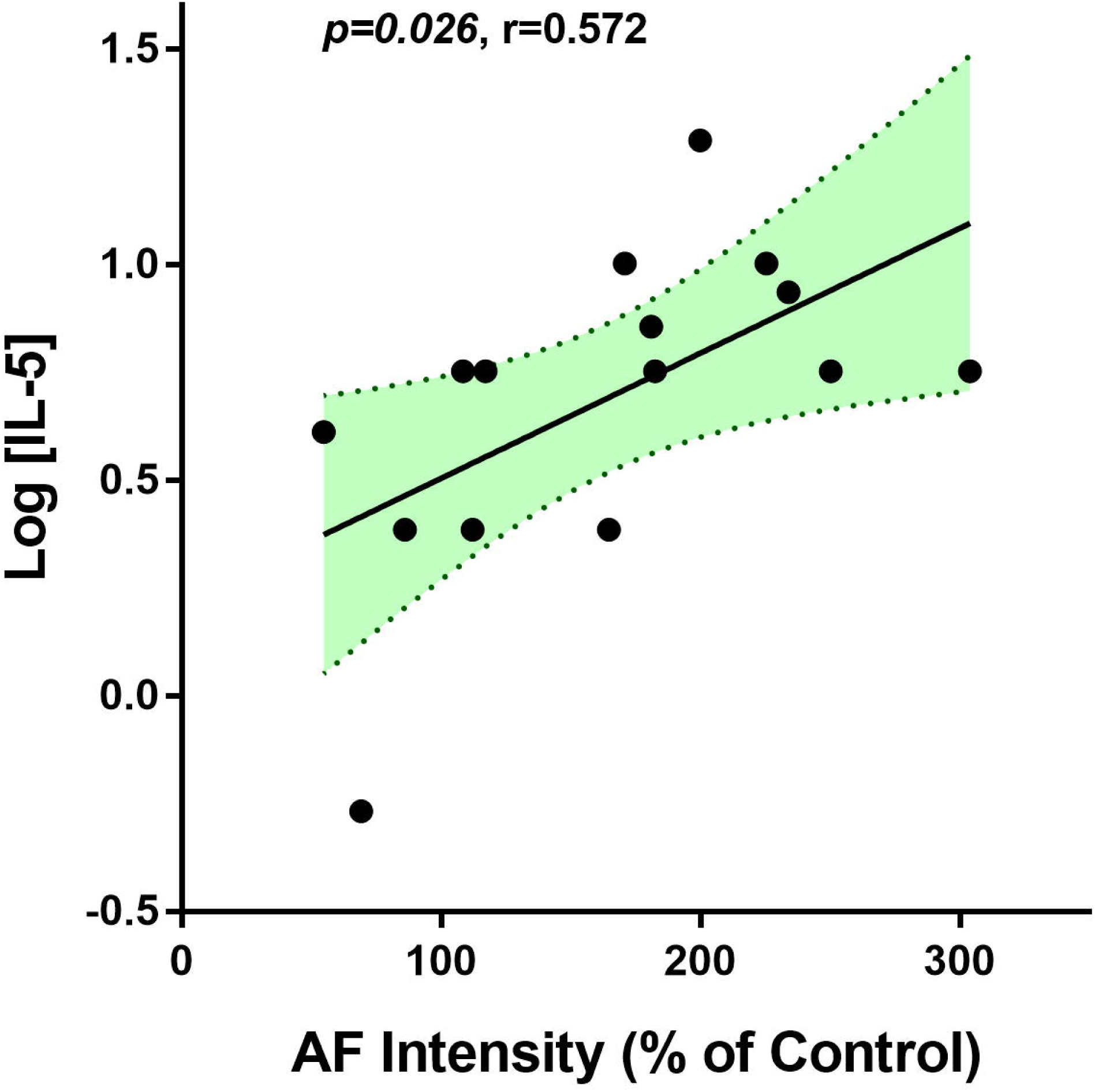
Both LPS doses and the epidermal green AF intensity were significantly associated the serum level of IL-5 in LPS-exposed C57BL/6Slac mice. Both LPS doses (A) and the epidermal green AF intensity (B) were significantly associated the serum level of IL-5 in LPS-exposed C57BL/6Slac mice. Each day the mice were i.p. administered with 0.5 or 1 mg / kg LPS. Three days after the first LPS administration, the serum level of IL-5 was determined. The serum IL-5 level (in pg/ml) was log-transformed. There were 4 – 6 mice in each group. The area in green shadow represents 95% confidence interval around the fitted line at the population level.

**Fig. 12.**
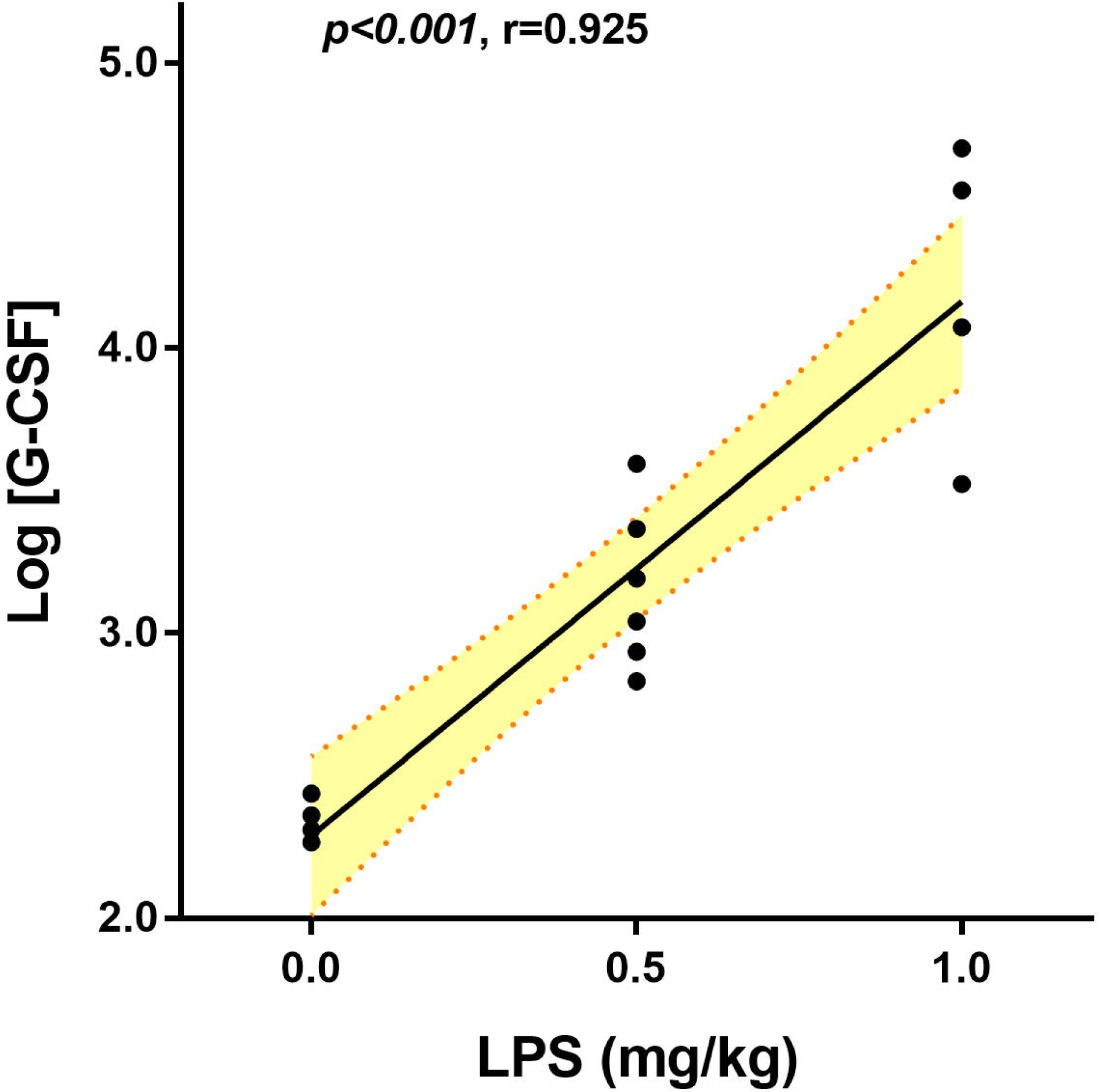
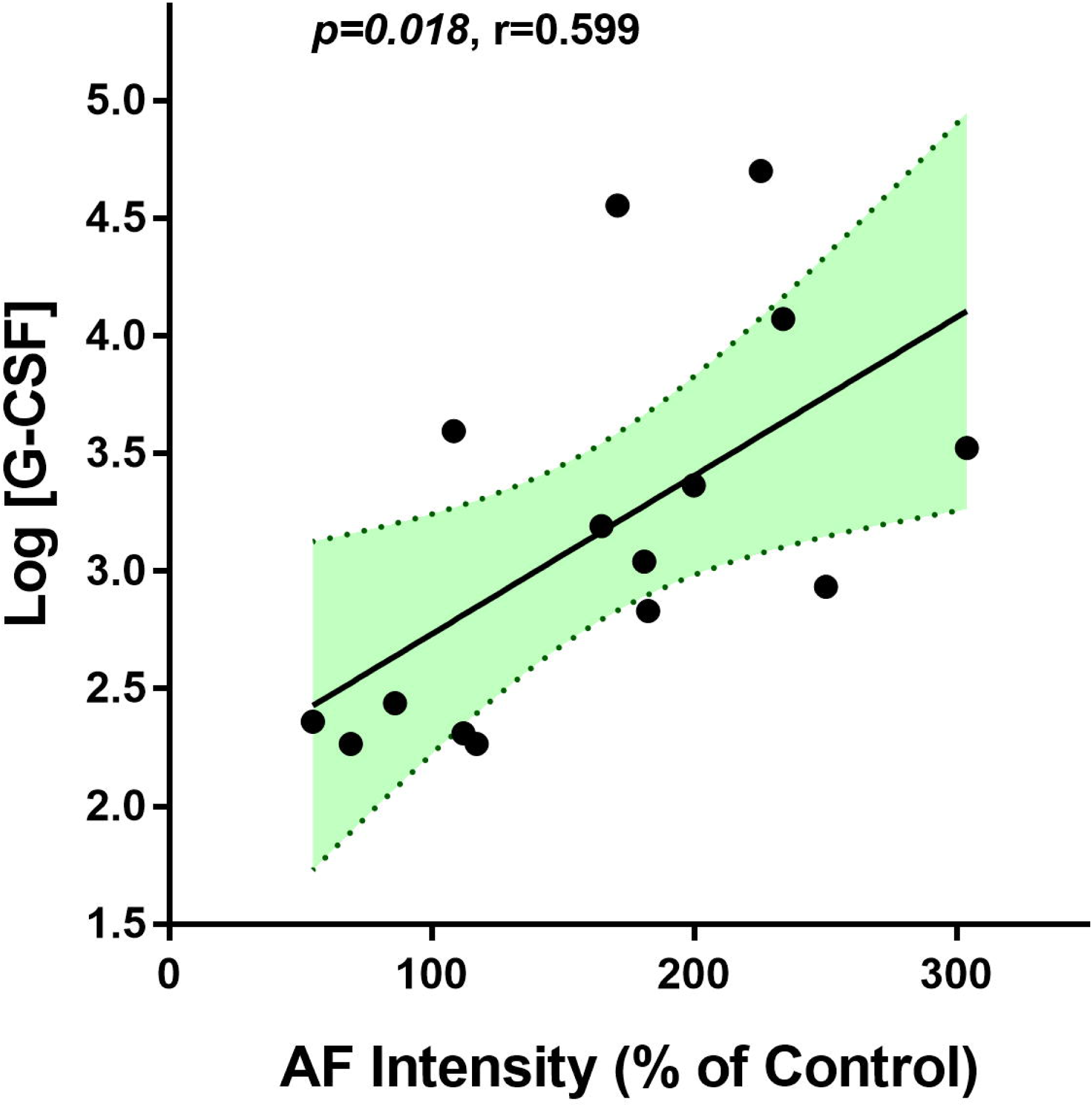
Both LPS doses and the epidermal green AF intensity were significantly associated the serum level of G-CSF in LPS-exposed C57BL/6Slac mice. Both LPS doses (A) and the epidermal green AF intensity (B) were significantly associated the serum level of G-CSF in LPS-exposed C57BL/6Slac mice. Each day the mice were i.p. administered with 0.5 or 1 mg / kg LPS. Three days after the first LPS administration, the serum level of G-CSF was determined. The serum G-CSF level (in pg/ml) was log-transformed. There were 4 – 6 mice in each group. The area in green shadow represents 95% confidence interval around the fitted line at the population level.

### 4. Neither LPS doses nor the epidermal green AF intensity were significantly associated with Log-transformed serum levels of several cytokines in LPS-exposed C57BL/6Slac mice

TNF-α is a cytokine that plays a key role in host defence against bacterial, viral and parasitic infection (3). Interferon-γ (IFN-γ) plays critical roles in both tumor control and the immunity against intracellular pathogens, while abnormal INF-γ levels are associated with multiple autoimmune and autoinflammatory diseases (25). Since none of the serum level of TNF-α, IFN-γ, IL-3, IL-4, IL-9,IL-12(p70), IL-13, IL-17A or granulocyte-macrophage colony-stimulating factor (GM-CSF) was in normal distribution, Log transformation was conducted for the data association analysis. Neither LPS doses nor the epidermal green AF intensity was significantly associated with Log-transformed serum level of these cytokines (Table 1).

**Table 1.** Neither LPS doses nor the epidermal green AF intensity was significantly associated with Log-transformed serum levels of several cytokines in LPS-exposed C57BL/6Slac mice. Each day the mice were i.p. administered with 0.5 or 1 mg / kg LPS. Three days after the first LPS administration, the serum level of TNF-α, IFN-γ, IL-3, IL-4, IL-9, IL-12(p70), IL-13, IL-17A or GM-CSF was determined. The serum cytokine levels (in pg/ml) were log-transformed. There were 4 – 6 mice in each group.

| Cytokine | [LPS] vs. Log [Cytokine] |  | AF vs. Log [Cytokine] |  |
| --- | --- | --- | --- | --- |
|  | *p value* | R | *p value* | R |
| IL-3 | 0.538 | 0.173 | 0.660 | 0.124 |
| IL-4 | 0.723 | 0.104 | 0.670 | 0.125 |
| IL-9 | 0.087 | 0.456 | 0.197 | 0.353 |
| IL-12(p70) | 0.879 | 0.043 | 0.847 | 0.055 |
| IL-13 | 0.050 | 0.514 | 0.279 | 0.299 |
| IL-17A | 0.311 | 0.281 | 0.441 | 0.215 |
| GM-CSF | 0.335 | 0.267 | 0.855 | 0.052 |
| IFN-γ | 0.085 | 0.459 | 0.177 | 0.368 |
| TNF-α | 0.246 | 0.320 | 0.432 | 0.219 |

Since both the serum Eotaxin concentrations and keratinocyte-derived chemokine (KC)/CXCL1 concentrations were in normal distribution, no Log transformation was needed for the data association analysis. Neither LPS doses nor the epidermal green AF intensity was significantly associated with the serum level of KC (Table 2). In contrast, LPS was not significantly associated with the serum level of Eotaxin, while epidermal green AF intensity was significantly associated with the serum level of Eotaxin (Table 2)

**Table 2.** LPS doses was not significantly associated with the serum levels of Eotaxin or KC. Neither LPS doses nor the epidermal green AF intensity was significantly associated with the serum level of KC. In contrast, LPS was not significantly associated with the serum level of Eotaxin, while epidermal green AF intensity was significantly associated with the serum level of Eotaxin. Each day the mice were i.p. administered with 0.5 or 1 mg / kg LPS. Three days after the first LPS administration, the serum level of Eotaxin or KC was determined. There were 4 – 6 mice in each group.

| Cytokine | [LPS] vs. [Cytokine] |  | AF vs. [Cytokine] |  |
| --- | --- | --- | --- | --- |
|  | *p value* | **R** | *p value* | **R** |
| **Eotaxin** | 0.291 | 0.292 | 0.016 | 0.611 |
| **KC** | 0.734 | 0.096 | 0.564 | 0.162 |

## Discussion

The major findings of our current study include: First, both the epidermal green AF intensity and LPS doses are significantly associated the serum levels of critical cytokines including IL-1β, IL-6 and IL-10 in LPS-exposed mice; second, both the epidermal green AF intensity and LPS doses are significantly associated the serum levels of pro-inflammatory factors including IL-2, IL-12(p40), MCP-1, MIP-1α, MIP-1β, and RANTES in LPS-exposed mice; and third, both the epidermal green AF intensity and LPS doses are significantly associated the serum levels of anti-inflammatory factors including IL-5 and G-CSF in LPS-exposed mice.

Inflammation is a key pathological factor in numerous diseases (15,19,23,24). Pro-inflammatory and anti-inflammatory cytokines are important mediators of inflammation (1-5,9,12,14,16,25,27). However, so far blood drawing is required for determining the cytokine levels in the blood, which constitutes a major obstacle for frequent monitoring of the inflammatory state of human subjects. It is of great scientific and clinical significance to search for non-invasive approaches for evaluating cytokine levels in the blood. Our previous finding has indicated that the epidermal green AF may become the first biomarker for non-invasive evaluations of systemic inflammation as well as numerous inflammation-associated diseases (17,29). However, it has remained unknown if epidermal green AF intensity is also significantly associated with the cytokines levels in the blood.

A key finding of our current study is that the serum levels of three widely studied cytokines, including IL-1β, IL-6 and IL-10, are significantly associated with the epidermal green AF intensity in the LPS-exposed mice. This finding has suggested that we may evaluate the levels of these important cytokines by determining non-invasively the AF intensity. Our study has also found that the AF intensity is not only associated with multiple pro-inflammatory factors including IL-2, IL-12(p40), MCP-1, MIP-1α, MIP-1β, and RANTES, but also two anti-inflammatory factors including IL-5 and G-CSF in LPS-exposed mice. Since our previous study has shown strong correlation between LPS doses and epidermal green AF intensity (17,29), the significant association between LPS doses and the serum levels of the cytokines could account for the significant association between the epidermal green AF intensity and the cytokines.

Based on our findings indicating the significant associations between the AF intensity and the serum levels of the cytokines, the AF intensity may become the first biomarker for non-invasive evaluations of serum levels of certain cytokines. The establishment of this non-invasive approach may profoundly enhance our capacity to monitor the health state, disease state and therapeutic effects of human subjects, since requirement of blood drawing and cost of the assays for inflammation have been the major obstacles for efficient and frequent monitoring. It is warranted to conduct studies to further establish solid approaches for determining the serum levels of certain cytokines, based on non-invasive detection of epidermal green AF intensity of human subjects. Establishment of our portable device for detection of epidermal green AF of human subjects (6,7,18,28) could significantly enhance our capacity to develop the non-invasive approaches.

Since the structure of the LPS-induced green AF is similar with the UV-induced green AF (13), which exhibits the characteristic polyhedral structure of the keratinocytes in *Stratum Spinosum*, the LPS-induced green AF could also be located in the keratinocytes in *Stratum Spinosum*. Because the only epidermal fluorophores that selectively exist in the keratinocytes of *Stratum Spinosum* are keratin 1 and keratin 10 (8,20,26), our study has suggested that keratin 1 and/or keratin 10 are the origin of the LPS-induced increases in the epidermal green AF. Future studies are needed to determine which cytokines mediate the alterations of the keratins to produce increased epidermal green AF in the LPS-exposed mice.

## Acknowledgment

The authors would like to acknowledge the financial support by a Major Special Program Grant of Shanghai Municipality (Grant # 2017SHZDZX01) (to W.Y.) and a Major Research Grant from the Scientific Committee of Shanghai Municipality #16JC1400500 and #16JC1400502 (to W.Y.).

